# Is a dam-altered river in the U.S. Southwest a barrier to dispersal for populations of a common lizard, *Uta stansburiana?*

**DOI:** 10.64898/2026.04.17.719235

**Authors:** Tessa C. Corsetti, Faith M. Walker, P. Brandon Holton, Daniel E. Sanchez, Gerard J. Allan, Jacque A. Lyman, Carol L. Chambers, Paul Beier

## Abstract

Dams can significantly alter natural riverine systems, but their impact on movement across rivers for most terrestrial vertebrates is poorly known. The completion of Glen Canyon and Flaming Gorge dams in Arizona and Utah (southwestern United States) profoundly changed the Colorado and Green Rivers and have altered habitat for many species. The common side-blotched lizard (*Uta stansburiana*) offers an excellent opportunity to examine the effects of riverine impoundments on migration and gene flow in terrestrial biodiversity. To assess these effects, we collected tissue samples from 241 *Uta stansburiana* above and below Glen Canyon Dam and on both sides of the Colorado river at three separate study areas. We used eight microsatellite loci to estimate genetic exchange in the context of genetic diversity and structure. One study area below Flaming Gorge Dam and above Glen Canyon Dam has annual periods of warmer water temperatures and lower flows that are closer to pre-dam conditions, whereas two study areas below Glen Canyon Dam have cold water temperatures year-round, and less pronounced seasonal low flow episodes. We predicted that warmer water temperatures above Glen Canyon Dam would promote greater genetic exchange among populations than below the dam. However, we found evidence for low levels of genetic exchange between sites both above and below Glen Canyon Dam, and a moderate amount of exchange at a site below this dam where lizards could conceivably move from one side to the other. Our results imply that 1) the changes in water temperature and hydrology in dam-altered rivers are a barrier for this species even when the distance from the dam is great; and 2) genetic exchange may be dependent on river morphology. These results are relevant to other small vertebrates, particularly ectotherms, that occupy habitat proximal to a dammed river and has implications for the conservation management of impounded river systems.

## Introduction

Free flowing rivers, those not significantly altered by dams or other anthropogenic influences, comprise only one third of global rivers [1, 2]. In the United States, just 25% of rivers remain free flowing [1]. Dam construction has immediate and long-term effects on aquatic and riparian systems [3, 4]. Although dam effects on vegetation, sediment flow, aquatic invertebrates/fishes, and natural river integrity have been widely studied, the effects of post-dam river hydrology as a potential barrier to migration and gene flow in terrestrial vertebrates, specifically reptiles, have been less well evaluated [2, 5, 6, 7]. Post-dam rivers limit genetic exchange between populations across taxa (e.g. riparian shrubs (*M. germanica)* in central Europe [3], platypus (*Ornithorhynchus anatinus)* in south-east Australia [4], and desert bighorn sheep (*Ovis canadensis nelson)* in the American southwest [6]), but there is little available research regarding how a dam-altered river may inhibit terrestrial reptiles.

In the absence of barriers, movement between populations can create meta-populations, allowing wildlife to recolonize patches after local extirpation, thereby sustaining the genetic diversity needed to counterbalance changing conditions and stochastic events [8]. Populations that lose connectivity due to fragmentation imposed by dams can experience a loss of genetic diversity and become less resilient to disease, severe weather events, and invasive species [8, 9]. The 1963 completion of Glen Canyon Dam on the Colorado River in the Grand Canyon (northern Arizona, USA) altered key riverine characteristics, with the 446 km stretch of river below the dam exhibiting a relatively constant flow rate, colder temperatures, and a loss of dynamic, seasonal flooding [7, 10]. These changes are particularly important in the context of how traversable the river has become for some species at certain times of the year [7, 11]. Large-bodied organisms may be able to counteract the barrier created by a dam-altered river by flying or strong swimming but smaller organisms, such as terrestrial lizards, may face an impenetrable barrier [12].

Studying dam effects on riverine systems is often complicated by a lack of pre-dam data. Historically, highly variable pre-dam flows carried an estimated 230,000 tons of sediment per day through the Grand Canyon, a large percentage (99%) of which is now trapped behind the dam. Flows below the dam have resulted in erosion of sandbars, eddies, and beaches through Grand Canyon [10, 13].

Above Glen Canyon Dam, the Colorado River through Cataract Canyon, Utah is the least modified stretch of the Colorado River mainstem [5] and experiences some flow regulation from Flaming Gorge Dam on the Green River, a major Colorado River tributary, in northern Utah. Flaming Gorge Dam was completed in 1962 and is ∼650 kms upriver of Cataract Canyon. The river through Cataract Canyon is more variable and the water temperature warmer. Although pre-dam temperature, sediment, and discharge data are limited, those recorded in the last decade for Cataract Canyon show water temperatures consistent with seasonal flooding. Median diurnal temperatures can be as low as 0°C during spring snowmelt and reach as high as 28°C during low flows in the late summer/ fall [14]. Water temperature below Glen Canyon Dam is now less variable, ranging from 8–15°C, with the median consistently at 10–11°C [14]. Sandbars occasionally emerge during low flows above the Glen Canyon Dam, but have been lost below the dam, and when combined with the lack of seasonal low flows, opportunities for terrestrial wildlife crossings are decreased [7, 13].

Due to the impacts imposed by Glen Canyon Dam, we hypothesized that there may be adverse effects on movement and gene flow of terrestrial ectotherms from one side of the Colorado River to the other. Specifically, we hypothesize that populations of the common side-blotched lizard (*Uta stansburiana*) below Glen Canyon Dam, which are exposed to consistent high-water discharge of low temperature, will have greater signatures of genetic structure than the population above the dam that, while influenced by Flaming Gorge Dam, is exposed to greater water fluctuations and higher temperatures. Accordingly, we assessed whether changes to river flow and temperature imposed by Glen Canyon Dam have altered both migration and genetic exchange in *Uta stansburiana. Uta stansburiana* is widespread in the American West and is abundant along the Colorado River corridor through the Grand Canyon [15, 16]. Therefore, this species may constitute an important test case for how reptiles of similar size and ecology can be affected by dam-altered rivers.

We examined genetic exchange in the context of genetic diversity and structure in *Uta stansburiana* at three study sites with two sampling areas, one on either side of the river, above and below Glen Canyon Dam during the summers of 2020 and 2021. The potential effect of a linear barrier, such as a highway or river, can be quantified by measuring the genetic diversity and structure of populations on either side [17, 18]. Assessment of genetic exchange (or lack thereof) requires a measure of genetic distance on each side of the barrier, and the putative barrier must have existed long enough to have influenced genetic composition in two or more populations [18]. Genetic divergence should be evident within 10–20 generations for populations consisting of 60-100 effective breeding individuals [16, 18]. Density of *Uta stansburiana* varies by time of year and predator activity but mean population density is ∼100 individuals per 2 ha site [19]. Mean generation time is ∼1 year [19], and the Glen Canyon Dam has existed for >55 years. If the cold, high flow below Glen Canyon Dam on the Colorado River is a barrier, its effect on genetic diversity and gene flow in *Uta stansburiana* should be evident today (while recognizing that genetic patterns were also influenced by historical population fluctuations).

## Materials and Methods

### Study areas and sampling sites

Our design involved sampling above Glen Canyon Dam where seasonal fluctuations in water flow and temperature resemble pre-dam conditions (Above-dam Site 1 = AD1). We also sampled two additional sites below the dam (Below-dam Site 1 = BD1; Below-dam Site 2 = BD2), where water flow and colder temperatures are more consistent (Fig 1). Sites were selected based on accessibility and available *Uta stansburiana* habitat on both the right and left sides of the river. We compared cross-river gene flow at one site above the dam (AD1) to two sites below the dam (BD1 and BD2).

**Fig 1.**
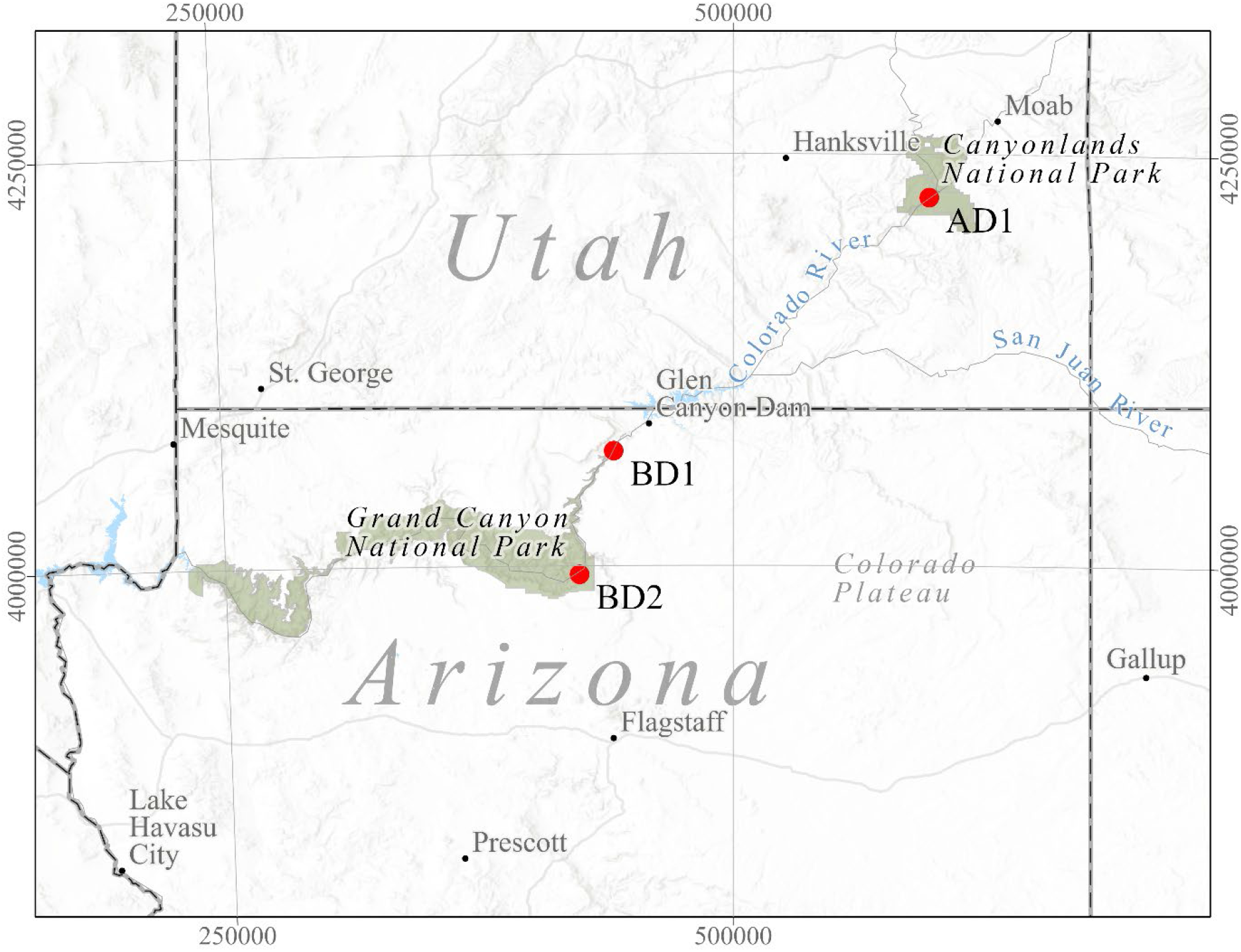
Research sites (filled red circles) on the Colorado River, Arizona and Utah. Above-dam Site 1 (AD1) in Cataract Canyon, Canyonlands National Park, Utah, and two sites below the dam in Grand Canyon National Park, Arizona: Below-dam Site 1 (BD1) and Below-dam Site 2 (BD2). Glen Canyon Dam is in Page, Arizona on the border of Arizona and Utah.

Although the above-dam cross-river sampling areas (S1 Fig) were potentially influenced by Flaming Gorge Dam on the Green River, they maintain approximate pre-dam river conditions and thus represent the least anthropogenically modified reach of the Colorado River mainstem [5, 7]. AD1 is primarily composed of sandy substrate with large boulders, gravel, and biocrust throughout [5]. It is dominated by native and non-native riparian desert scrub species such as saltcedar (*Tamarix spp*.), Russian thistle (*Salsola spp*.), brome grasses (*Bromus spp*.), prickly pear (*Opuntia spp*.), and cottonwood *(Populus fremontii)* [5].

BD1 is 37 km downriver from Glen Canyon Dam (S2 Fig). BD1.R (river right) and BD1.L (river left) are within dry side canyons occasionally scoured by seasonal flooding. They are primarily composed of sandy substrate interspersed with large boulders, gravel, and biocrust, and contain sparse vegetation dominated by saltcedar (*Tamarix spp*.), Mormon tea *(Ephedra nevadensis)*, shadescale *(Atriplex confertifolia)*, and prickly pear (*Opuntia spp*.) [20].

BD2 (S3 Fig) is 135 km downriver from Glen Canyon Dam. Both BD2.L (river left) and BD2.R (river right) are large, sandy sites with large boulders and gravel throughout. BD2.R is ∼1km downriver from BD2.L and has an abundance of biocrust amidst sandy, brushy substrate. Vegetation is primarily desert scrub with saltcedar (*Tamarix spp*.), brome grasses (*Bromus spp*.), prickly pear (*Opuntia spp*.), Mormon tea *(Ephedra nevadensis)*, catclaw acacia *(Acacia greggii)*, and common reed *(Phragmites australis)* [20]. The river morphology between BD2.L and BD2.R is less linear than other study areas, with side canyon debris flows creating an abrupt, meandering effect (S3 Fig).

### Sample Collection

*Uta stansburiana* is a small (<64 mm snout-vent length) lizard with both sexes easily identified by a prominent blue-black spot posterior to its front legs [16]. It is primarily found in sandy, desert scrub vegetation. Home range averages ∼446 m^2^ (maximum 810 m^2^) for males and ∼121 m^2^ (maximum 202 m^2^) for females, with individual lizards defending small-area microhabitats within a larger home range [16, 19]. *Uta stansburiana* expands its home range directly in proportion to age, with hatchlings remaining in proximity of their hatch location for several weeks [16]. Dispersal distance is influenced by habitat, time of hatching, and sex [21]. Both sexes exhibit elevated site fidelity, are highly territorial, and are most often observed on the ground or sunning themselves on rocks, rarely climbing into trees or shrubs unless pursued or threatened [16, 19]. Widespread in the western United States, *Uta stansburiana* ranges from central Washington to the tip of Baja California and from the Pacific coast to western Colorado and western Texas [16]. The species is currently stable across its range and is designated as being of least concern on the IUCN Red List of Threatened Species [22].

We sampled *Uta stansburiana* along shoreline habitat and within side canyons perpendicular to the river until a minimum sample size of 25–30 individuals were obtained [23, 24]. We caught adult lizards with an unwaxed dental floss lasso. We recorded sex, approximate capture and release time, and tail condition, then clipped 5-7 mm of tissue from the tip of the tail. Samples were stored in RNA*later* (Invitrogen, Thermo Fisher Scientific, Waltham, MA, USA), then stored at -20℃ prior to DNA extraction. Lizards were marked with a non-toxic xylene-free paint marker in colors consistent with their natural coloration to prevent recapture and resampling [25] and released at point of capture.

### DNA Extraction and Standardization

We performed DNA extractions using Qiagen’s DNeasy Blood & Tissue Kits (Qiagen, Germantown, Maryland, USA) according to manufacturer’s protocol. Tissue samples were lysed overnight at 55°C in the provided lysis buffer, with final elution at 200 µL. DNA extracts were quantified on a Nanodrop 8000 Spectrophotometer (Thermo Fisher Scientific, Waltham, MA, USA), and standardized to 5 ng/µL using molecular grade water. Samples with starting concentrations lower than 5 ng/µL were kept at their original concentrations.

### Microsatellite Loci and PCR Conditions

We adapted nine microsatellite primers with a universal tail labeling system [26, 27]. We PCR-amplified all loci in reactions containing 1X PCR buffer (Invitrogen, Thermo Fisher Scientific, Waltham, MA, USA), 2 mM of MgCl2, 0.2 mM dNTPs, 0.1 mM labeled forward primer, 0.2 mM unlabeled reverse primer, 0.2 mM fluorescent label, 0.08 U/µL Platinum Taq DNA polymerase (Invitrogen), 0.02 µg/µL Ultrapure non-acetylated Bovine Serum Albumin (BSA), and 5 µL of DNA template in a 15 µL reaction [28]. We performed all thermal cycling on an Applied Biosystems SimpliAmp Thermal Cycler. Each reaction followed modified conditions from Alloush 2013 [29] beginning with 90°C for 2 min, followed by 45 cycles of 95°C for 1 min, 51°C for 1 min, and 72°C for 1 min, then concluded with a final extension of 72°C for 5 min. Following amplification, PCR products were diluted as follows with molecular grade water prior to fragment analysis on an Applied Biosystems 3130 Genetic Analyzer: 1:100 for loci 10,000 M’s, SMcL, and SVeg; 1:10 for loci BRtt, IGs, MCC, NGff, PKln, and SPhill. Alleles and associated fragment sizes were scored using GeneMarker software (SoftGenetics, State College, PA, USA).

### Quality Control

To minimize genotyping errors, we adjusted for scoring errors, null alleles, and PCR artifacts by repeating PCRs for novel alleles, poorly amplifying samples, and multi-locus genotypes that differed at three or fewer loci. We used Micro-Checker [30] to further assess scoring errors, allelic dropout, and null alleles. We utilized GenAlEx 6.5 [31] to test for linkage disequilibrium and departures from Hardy-Weinburg equilibrium (HWE).

### Genetic Analysis

Genetic diversity was represented by mean observed and expected heterozygosity and allelic richness. Rarefaction to correct unbalanced sample sizes was performed using the program HP-Rare [32]. We used Analysis of Molecular Variance (AMOVA; GenAlEx 6.5 [31]) to examine how genetic variation is partitioned between sampling areas on river right and river left across the three study sites. We used the fixation index (F_ST_) in AMOVA to measure population differentiation due to genetic structure (GenAlEx 6.5) and adjusted for multiple tests with sequential Bonferroni correction. We performed a Principal Coordinates Analysis (PCoA) in GenAlEx 6.5 to visualize the genetic distance between populations and then used discriminant analysis of principal components (DAPC) [33] to produce a graphical summary that maximized among-group variation and minimized within-group variation in our a priori defined populations [34].

We conducted two DAPC runs using R package adegenet in R 4.0.5 [35, 36]. The first run incorporated all populations while the second run was to visualize finer-scale differentiation between BD2.L and BD2.R. We built initial models for each run, retained 40 PCs for the first run and 25 for the second. To prevent over-fitting, we used the initial models to select the optimal number of PCs to retain (function: optim.a.score). We then built a final DAPC by retaining the optimal number of PCs given by the alpha-score result.

We used STRUCTURE [37] to determine de novo population genetic structure and estimate distinct populations within our dataset. We used 20 runs per K (K=1-8), a burn-in of 100,000, with 500,000 Markov Chain Monte Carlo (MCMC) iterations [38]. Results for each K were averaged and graphed using CLUMPAK [39], with genetic clusters (K) identified using Best K [37, 40].

To determine if there were parent-offspring pairs on opposite sides of the river, we used the log-likelihood-based program CERVUS 2.0 [41], with each individual (candidate offspring) tested with all candidate mothers and fathers, using separate simulations of 10,000 cycles to infer parentage. We used settings of 50% of candidate parents sampled, 97% of loci genotyped, and an error rate of 0.5%. Parentage was assigned when: 1) LOD > 3.0, 2) delta > 95%, and 3) no more than one allele mismatched by not more than two base pairs.

We evaluated directionality of recent gene flow (within ∼ 4 generations) using BAYESASS 3.0.4 [42]. Three separately seeded runs were conducted with 21,000,000 MCMC iterations, discarding the first 5,000,000 as burn-in, and sampling every 1,000 iterations (Mixing parameters: --deltaA = 0.3, --deltaF = 0.4, --deltaM = 0.2). We then checked for chain convergence among runs using Tracer 1.7.2 [43] and calculated 95% credible intervals following user manual recommendation.

## Results

We generated multilocus microsatellite data (n = 8 loci) for 241 unique *Uta stansburiana* across six putative populations in Canyonlands (AD1) and Grand Canyon (BD1, BD2) National Parks (Table 1). We were unable to collect the minimum sample size at two (of six) sampling areas (AD1.R and BD2.R).

**Table 1.** Research site and sample collection data for the 2020 and 2021 field seasons.

| Site | UTM | Location | River Side | # Samples |
| --- | --- | --- | --- | --- |
| AD1.R | 12 S 593457 E 4223754 | Canyonlands NP, Utah | Right | 20 |
| AD1.L | 12 S 594067 E 4223839 | Canyonlands NP, Utah | Left | 49 |
| BD1.R | 12 S 441337 E 4069939 | Grand Canyon NP, Arizona | Right | 64 |
| BD1.L | 12 S 441563 E 4069818 | Grand Canyon NP, Arizona | Left | 58 |
| BD2.R | 12 S 424351 E 3994857 | Grand Canyon NP, Arizona | Right | 16 |
| BD2.L | 12 S 424924 E 3995651 | Grand Canyon NP, Arizona | Left | 34 |

Research site and sample collection data including site name, UTM (NAD27/UTM Zone 12N), site location, river side (right or left), and the number of individuals collected at each research site.

### Genotyping

We excluded marker NGff from the original nine locus panel after scoring attempts failed to produce genotypes at the expected allele size for most samples. All eight loci were successfully genotyped for 97.5% of individuals, with failures non-specific to population or locus. One individual was genetically identified as a recapture and thus removed from the analysis. All eight loci were polymorphic (percent polymorphic loci across populations = 83.33%; SE = 4.17%) and showed no deviations from HWE nor evidence of linkage disequilibrium. Micro-Checker [30] confirmed no evidence of scoring errors, dropout of larger alleles, or null alleles.

Following rarefaction, BD2.L had the greatest allelic richness, followed by BD1.R, AD1.L, and BD1.L. Sites AD1.R. and BD2.R had the lowest allelic richness, potentially due to small sample sizes at both locations (N=20 and 16, respectively). Private alleles were detected in all populations (Table 2).

**Table 2.** Population genetic parameters of *Uta stansburiana*.

| Site | Mean<br>Individuals<br>per locus | Alleles per<br>Locus ± SE | Ho ± SE | He ± SE | Rarefied<br>Allelic<br>Richness | Private<br>Alleles |
| --- | --- | --- | --- | --- | --- | --- |
| AD1.R | 20 | 4.1 ± 1.0 | 0.49 ± 0.14 | 0.47 ± 0.13 | 3.9 | 1 |
| AD1.L | 48.9 | 5.0 ± 1.2 | 0.45 ± 0.12 | 0.43 ± 0.12 | 4.1 | 2 |
| BD1.R | 63.5 | 5.1 ± 1.0 | 0.48 ± 0.11 | 0.50 ± 0.11 | 4.1 | 3 |
| BD1.L | 57.8 | 4.6 ± 0.9 | 0.49 ± 0.11 | 0.52 ± 0.11 | 4.1 | 4 |
| BD2.R | 16 | 3.6 ± 1.0 | 0.41 ± 0.13 | 0.38 ± 0.12 | 3.6 | 2 |
| BD2.L | 34 | 4.8 ± 1.2 | 0.38 ± 0.12 | 0.40 ± 0.13 | 4.2 | 3 |

Population genetic parameters of *Uta stansburiana* in Grand Canyon and Canyonlands National Parks, USA. Includes mean number of individuals genotyped at each locus, the average number of alleles per locus, observed heterozygosity (Ho), expected heterozygosity (He), rarefied allelic richness, and the number of private alleles observed in each population.

### Cross-river Site Analyses

We found significant genetic structure between AD1.R and AD1.L (F_ST_ = 0.126, α = 0.003, p = 0.001; S4 Fig; Table 3). All pairwise F_ST_ estimates were significant for both permutation and bootstrapping tests. We also found significant genetic structure between BD1.L and BD1.R (F_ST_ = 0.124, p = 0.001; S5 Fig; Table 3), as well as between BD2.L and BD2.R (F_ST_ = 0.053, p = 0.001; S6 Fig; Table 3). Clumpak Best K analysis for probability by K (40; 43) grouped a core set of 50 individuals into a single breeding population. Estimations of populations of *Uta stansburiana* at BD2.L and BD2.R, Grand Canyon NP derived Δk for k from 1 to 4. There was one assumed population or genetic group between two sites, reflecting a small sample size at BD2.R or indicating that lizards can access and successfully breed with individuals on the opposite side of the river. Evanno et al. (2005) [40] determined Delta K=2, indicating two assumed populations or genetic groups in this study. Pritchard et al. (2000) [37] determined K=1, indicating one single population between these two sites.

**Table 3.** Pairwise Population F_ST_ Values (lower diagonal) between river right and river left populations in Canyonlands and Grand Canyon National Parks. Asterisks in the upper diagonal indicate significant P-values (α = 0.05). Bolded F_ST_ values show population differentiation between cross-river study sites.

| Pairwise Population $F_{ST}$ Values | | | | | | |
| --- | --- | --- | --- | --- | --- | --- |
|  | BD1.R | BD1.L | BD2.R | BD2.L | AD1.R | AD1.L |
| BD1.R | 0 | * | * | * | * | * |
| BD1.L | **0.124** | 0 | * | * | * | * |
| BD2.R | 0.234 | 0.237 | 0 | * | * | * |
| BD2.L | 0.221 | 0.249 | **0.053** | 0 | * | * |
| AD1.R | 0.143 | 0.182 | 0.146 | 0.158 | 0 | * |
| AD1.L | 0.205 | 0.245 | 0.25 | 0.255 | **0.126** | 0 |

The DAPC for all sites (9 PCs retained; Fig 2) demonstrated that above- and below-dam sites were well differentiated from one another. Save for BD1.L and BD2.R, no significant differentiation was noted between populations on both sides of the river at a given site.

**Fig 2.**
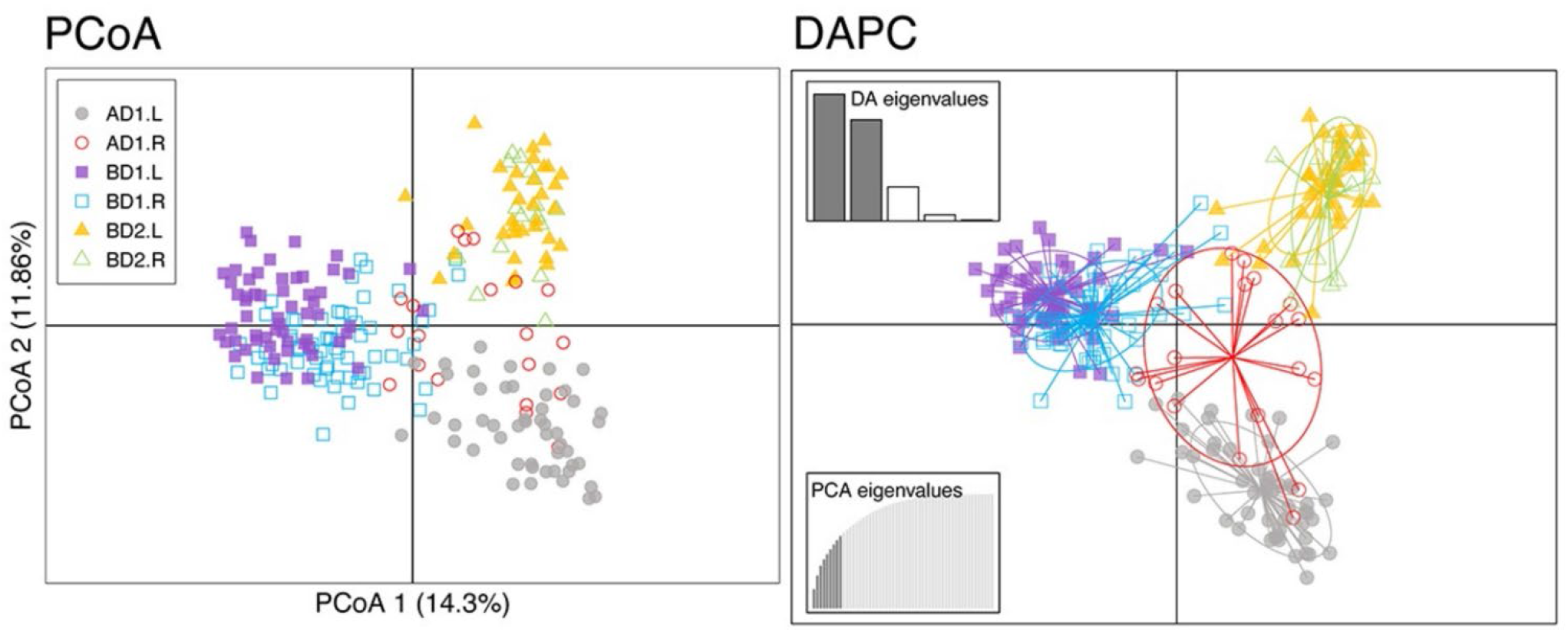
Principal Coordinates Analysis (PCoA) and a priori discriminant analysis of principal components (DAPC) for populations of *Uta stansburiana* above (AD) and below (BD) Glen Canyon Dam, AZ. The left panel shows a PCoA scatter plot of differentiation among all populations, isolation by distance shown using a pairwise genetic distance matrix. Analysis shows clustering of populations consistent with their geographic location. Inertia ellipses bound population centroids. The right panel shows a separate DAPC (represented as density plots) for the two BD2 sites to visualize their finer-scale differentiation. Samples are color-coded by population. River right sites are open symbols and river left sites are filled symbols.

All three separately seeded BAYESASS runs (S2 Table) exhibited MCMC chain convergence and produced identical estimates. A significant but asymmetric migration rate (95% CI did not overlap zero) was observed below the dam at river left (BD2.L) and river right (BD2.R) suggesting a modest amount of migration across the river. We observed no other significant migration rates among other sites and river sides.

### Analyses among all sites

We found significant levels of genetic structure (p = 0.001; Fig 2) among populations on river right vs. river left. Overall, this suggests that populations on the left and right sides of the river are genetically differentiated from one another, despite the larger distance between populations on the same side of the river.

K-values derived from one to eight potential groups [40] separated 241 *Uta stansburiana* into two populations, while STRUCTURE [37] grouped individuals into six populations. We identified substantial genetic structure in four (of six) populations (Fig 3). While K=6 was selected based on STRUCTURE analysis, K=5 provided a more stable structure, and thus both were included in our results. Parentage analysis revealed parent-offspring pairs on the same side of the river with 80% to 95% confidence, and no parent-offspring pairs on opposite sides of the river at any location (S3 Table).

**Fig 3.**
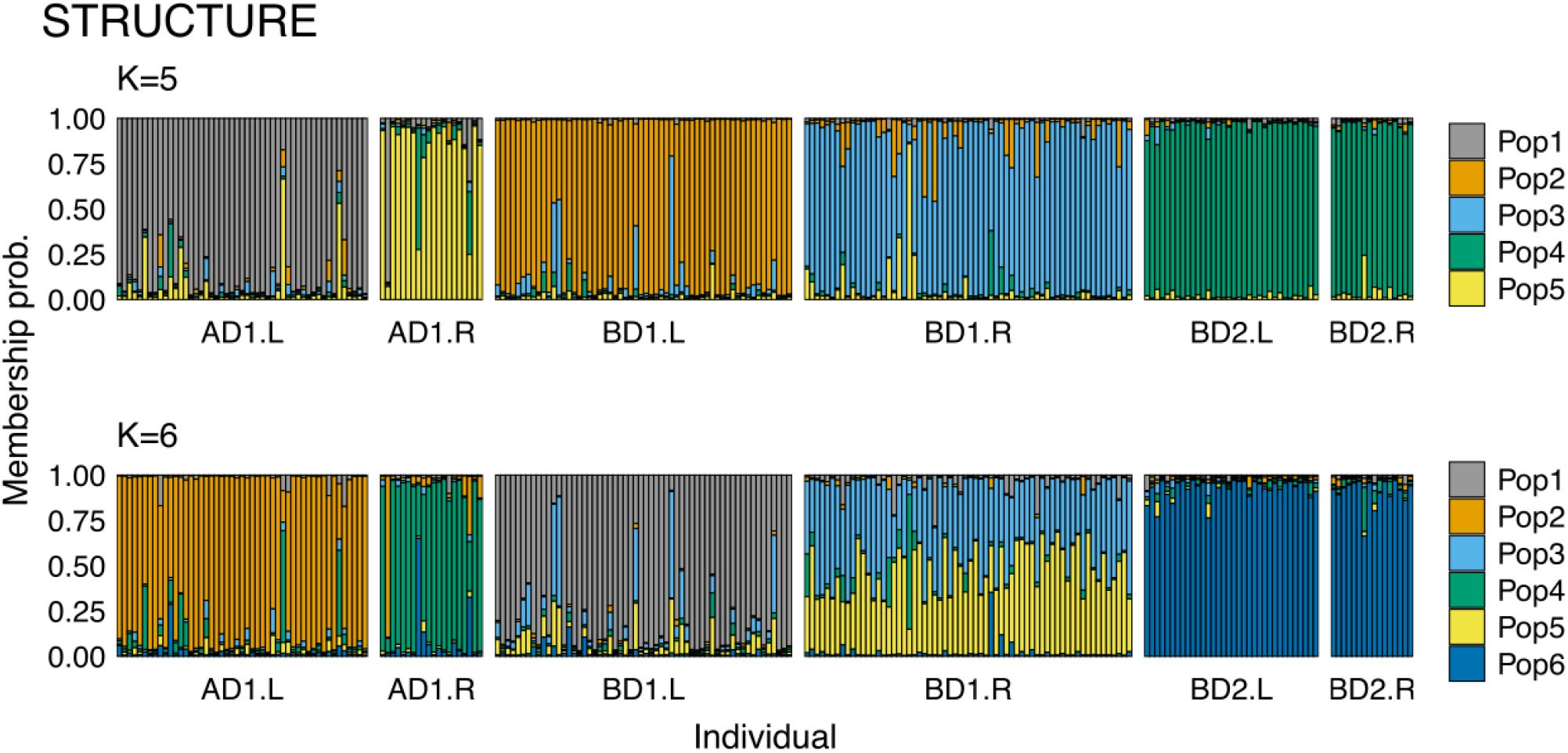
STRUCTURE for all *Uta stansburiana* populations. STRUCTURE result based on six populations (K=6) in Canyonlands and Grand Canyon National Parks. Each individual sample is represented by a vertical bar with colors indicating the proportions of each genetic cluster. Bar plot shows genetic structure with some admixture between BD1.R (1, blue) and BD1.L (2, red) and AD1.R (5, pink) and AD1.L (6, yellow). Populations BD2.L (3, teal) and BD2.R (4, green) are of mixed ancestry, suggesting a single breeding population.

## Discussion

Migration and geneflow are essential for the longevity and viability of natural populations and processes [3, 4, 18]. There is growing concern over the adverse effects that impoundments such as dams may have on movement and gene flow [4, 6]. Werth et al. [3] found strong effects of genetic isolation in populations of specialist riparian plant species in dam-altered riverine system in Central Europe. Mijangos et al. [4] determined that genetic differentiation in platypuses increased proportionately with time after a dam was built and was higher in dammed rivers than free-flowing rivers. Pirani et al. [48] established that rivers act as barriers to gene flow for Amazonian herpetofauna and stressed the importance of understanding the many processes associated with rivers as potential barriers as riverine systems continue to face pressures from climate change and human development.

Our analyses are consistent with the results reported by Pirani et al., Mijangos et al. and Werth et al. and suggest that the dam-altered Colorado River causes a disruption of movement and gene flow above in addition to below Glen Canyon Dam, indicating that even flows and temperatures moderated by distance from a dam (i.e., Flaming Gorge Dam) are sufficient for impeded dispersal. We found significant structure in two of the three study areas (AD1 and BD1) and populations clustered into regional groupings with overlap of BD2.L and BD2.R populations. The presence of private alleles at all sites suggested either limited gene flow between populations or high mutation rates of individual loci that exceed gene flow and migration [44].

STRUCTURE analysis indicated six populations with clear structure between BD1.R/BD1.L and AD1.R/AD1.L, suggesting limited migration and associated gene flow among these populations. Furthermore, parentage analysis showed no parent-offspring pairs on opposite sides of the river. Together, these data support our hypothesis that the river may act as barrier to movement and associated gene flow.

Genetic differentiation was observed separating AD1 and BD1 across the river, but we were surprised by the lack of separation between the BD2.L and BD2.R. We offer two possible hypotheses: First, lizards could be swept downstream from BD2.L and wash ashore at BD2.R, surviving the short river distance between the two sites, and augmented by the sites’ non-linear river morphology (S3 Fig). We encountered two lizards within the river during field work, one floating ∼1m from the shore alive but unable to swim at BD1.L, while the second was dead ∼1.5 m from shore at BD1.R. The occurrence of a live lizard in the water suggests they may survive being swept down river to wash up on the opposite shore, as the non-linear river morphology between BD2.L and BD2.R is vastly different from that separating other sampling areas. It is hypothesized that as few as one effective migrant every two generations is sufficient to maintain heterozygosity whereas five or more per generation would result in little to no genetic differentiation, although this is also contingent upon effective population size [9, 45]. If this hypothesis is applied, then a single lizard successfully traversing from BD2.L to BD2.R every two years could potentially maintain connectivity between the two populations. However, additional sites would need to be studied to determine whether a particular river morphology facilitates migration.

Our second explanation for low genetic divergence between BD2.L and BD2.R is the smaller sample size (N=16) at BD2.R [23]. Since allelic richness and private alleles are influenced by sample size, it is possible that the low genetic differentiation is an artifact of sampling, where private alleles may have been undetected [32]. Our K = 2 findings [40] should be carefully considered as potential exists for alternative groupings with increased sampling of BD2.L and BD2.R [46, 47].

Future studies would benefit from the assessment of other small vertebrates with life histories and migration patterns similar to *Uta stansburiana* (e.g., other small reptiles or amphibians). We also suggest that future studies would benefit from examining other types of molecular markers such as mitochondrial DNA, which due to its different pattern of inheritance (maternal vs. bi-parental) may reveal distinct haplotypes above and below the dam [17]. These data could provide important insight on how a maternally inherited marker is distributed above and below the dam, and in corresponding populations on either side of the river, leading to a better understanding of how dam-altered rivers affect migration and gene flow in different species. Finally, for our study system, it would be worthwhile to sample additional sites above and below Glen Canyon Dam, as well as sites along a free-flowing river, such as the Yampa River in northwestern Colorado, USA.

The Colorado River is one of the most manipulated and regulated bodies of water in the United States, with over a dozen large dams and thousands of small impoundments between its headwaters in the southern Rocky Mountains of Colorado to its historical endpoint in the Gulf of California [11]. Impoundments, dams, and irrigation have changed its historic sediment transport, flood dynamics, water temperature/ chemistry, hydraulic flow, and the distribution and abundance of native biota [10, 11]. Understanding how changes to river hydrology has affected wildlife will be beneficial for predicting potential effects of human development and climate change on riverine systems.

The limited ability for *Uta stansburiana* to migrate across the river in most cases, in combination with the genetic structure we observed among the study areas, suggests that if habitat degradation continues then small-bodied lizards constrained to isolated patches of favorable habitat may suffer loss of diversity within and between populations. While *Uta stansburiana* is common and widespread, dam-related changes in flow and temperature could have more profound effects on rare or endemic species. As natural processes continue to be disturbed by climate change and human development, identifying landscape characteristics, such as dam-altered rivers, that could be disrupting gene-flow and migration is a vital part of landscape and conservation management [46, 48].

## Acknowledgements

Research was conducted under National Park Service research permits for Grand Canyon National Park (No. GRCA-2020-SCI-0013) and Canyonlands National Park (No. CANY-2020-SCI-0010). Lizard capture and handling methods were approved through the National Park Service and Northern Arizona University Institutional Animal Care and Use Committees (Protocol #20-001).

We would like to thank Andrew Holycross for his valuable insight and guidance for this work, the Colorado Plateau Cooperative Ecosystem Studies Unit for their support, the many field technicians who volunteered their time to assist with sample collection, and National Park Service River rangers for their assistance in accessing field sites in the Grand Canyon and Canyonlands National Parks.

