## Supplementary figures and images for "Is a dam-altered river in the U.S. Southwest a barrier to dispersal for populations of a common lizard, *Uta stansburiana?*"

### Supplemental Figure 1

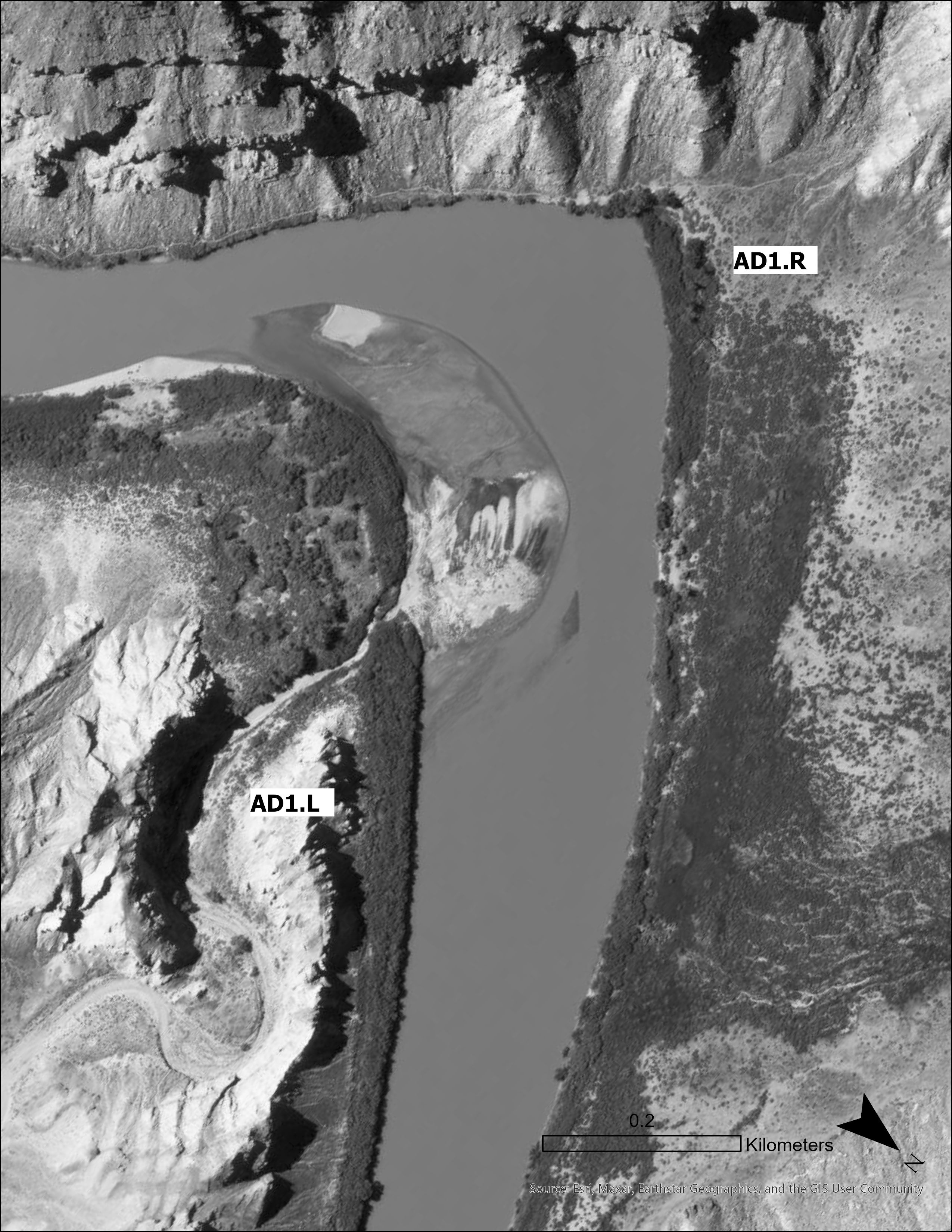

### Supplemental Figure 2

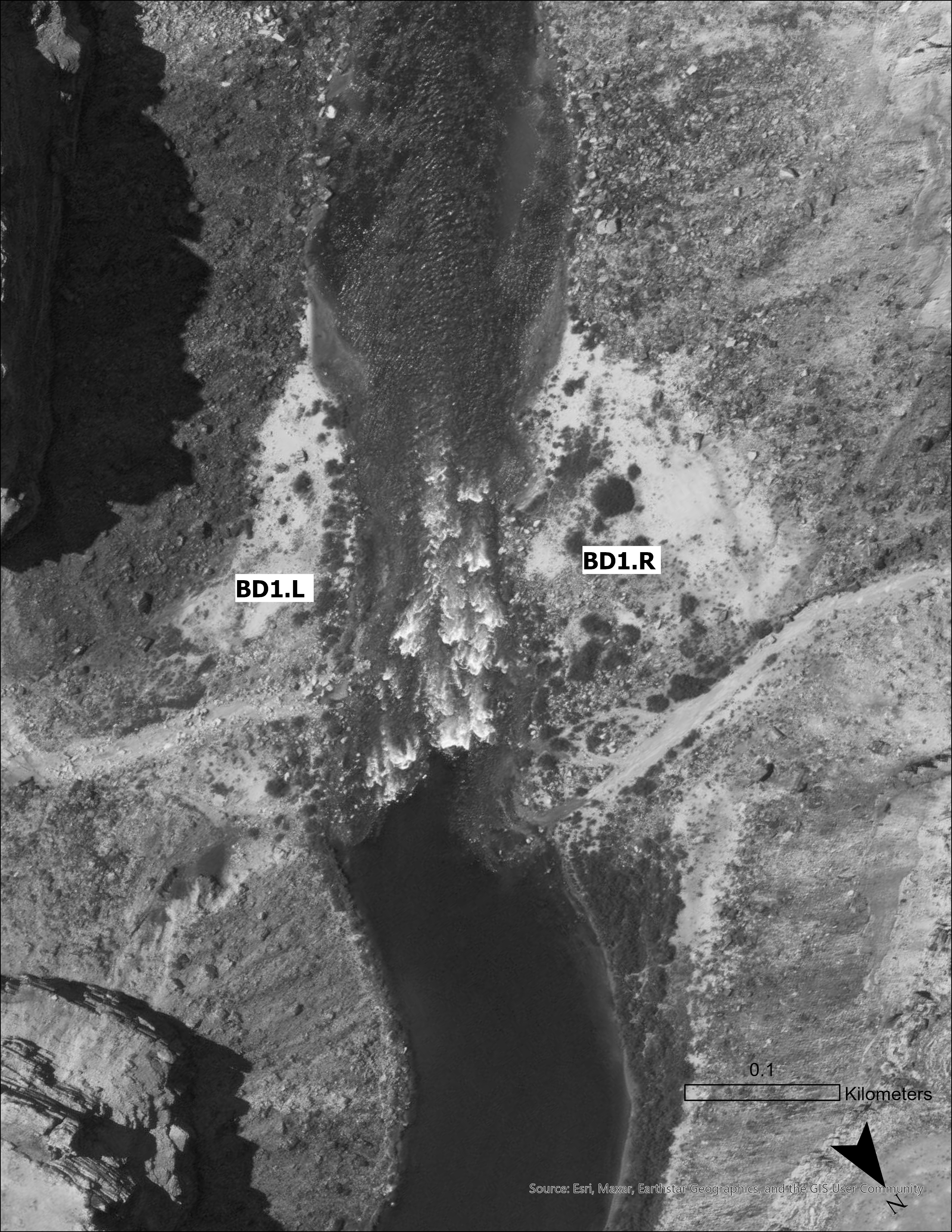

### Supplemental Figure 3

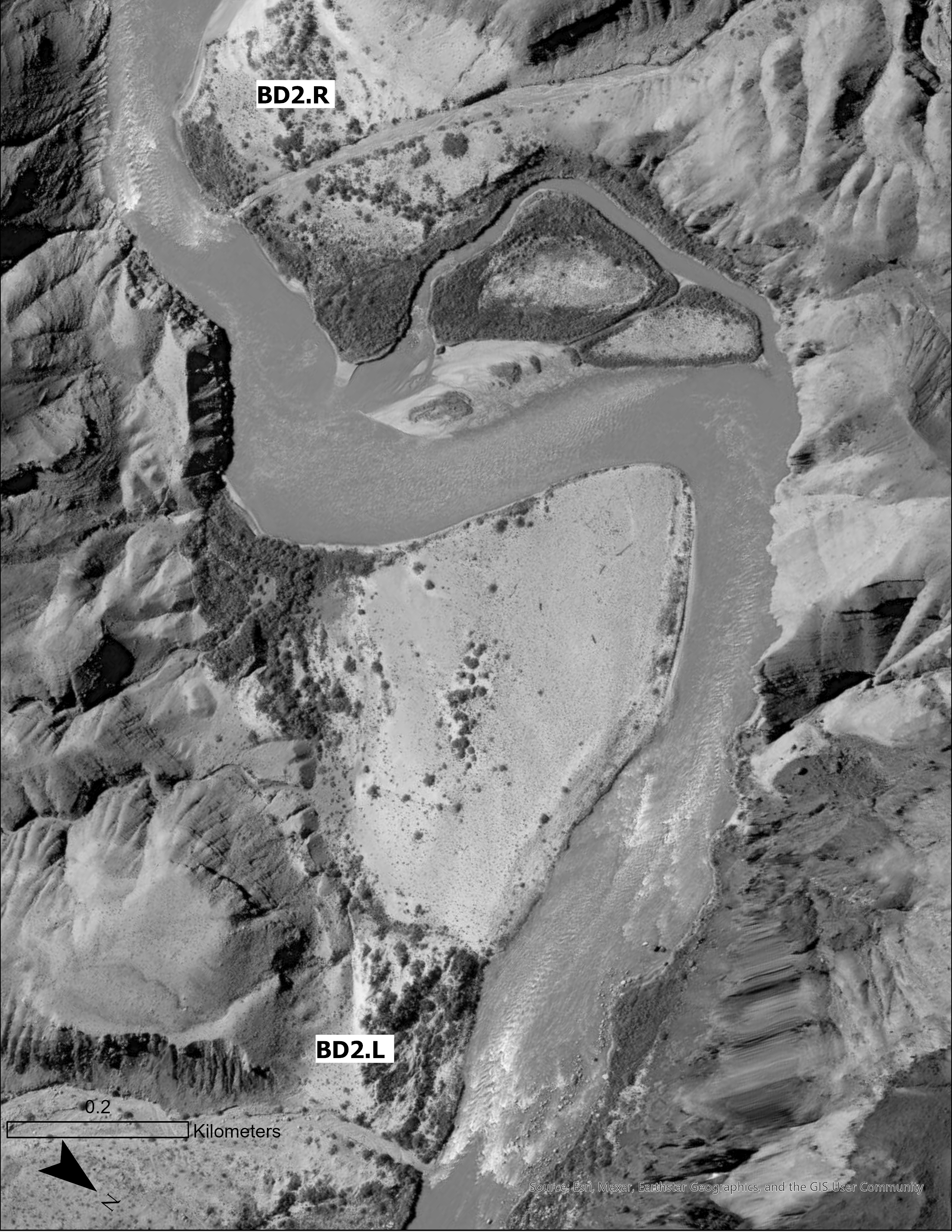

### Supplemental Figure 4A

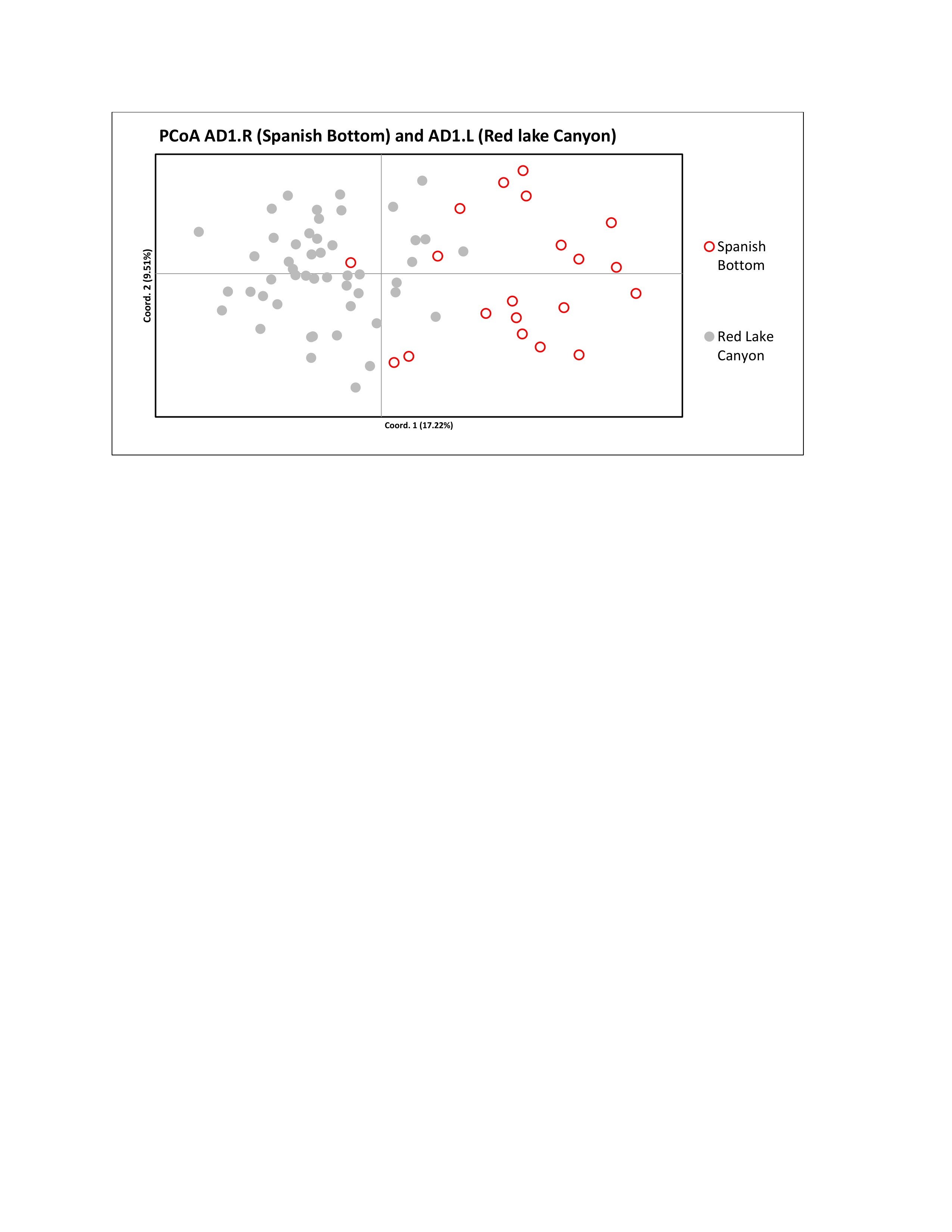

### Supplemental Figure 4B

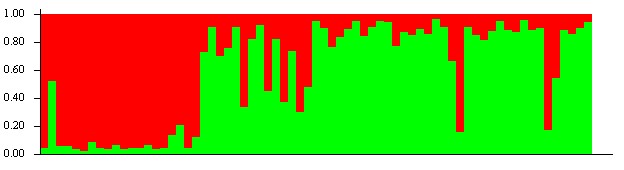

### Supplemental Figure 5A

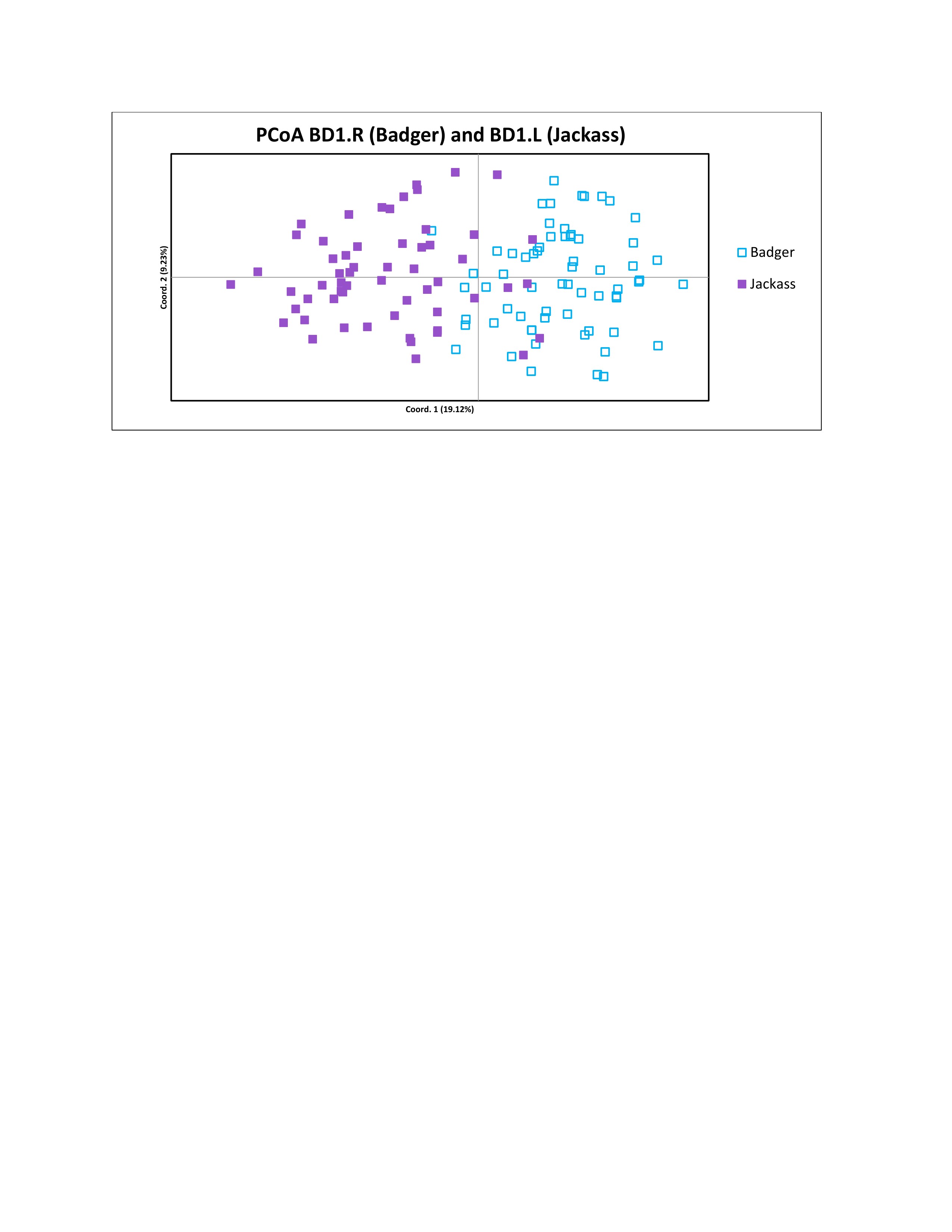

### Supplemental Figure 5B

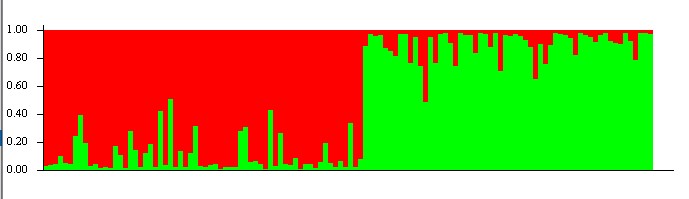

### Supplemental Figure 6A

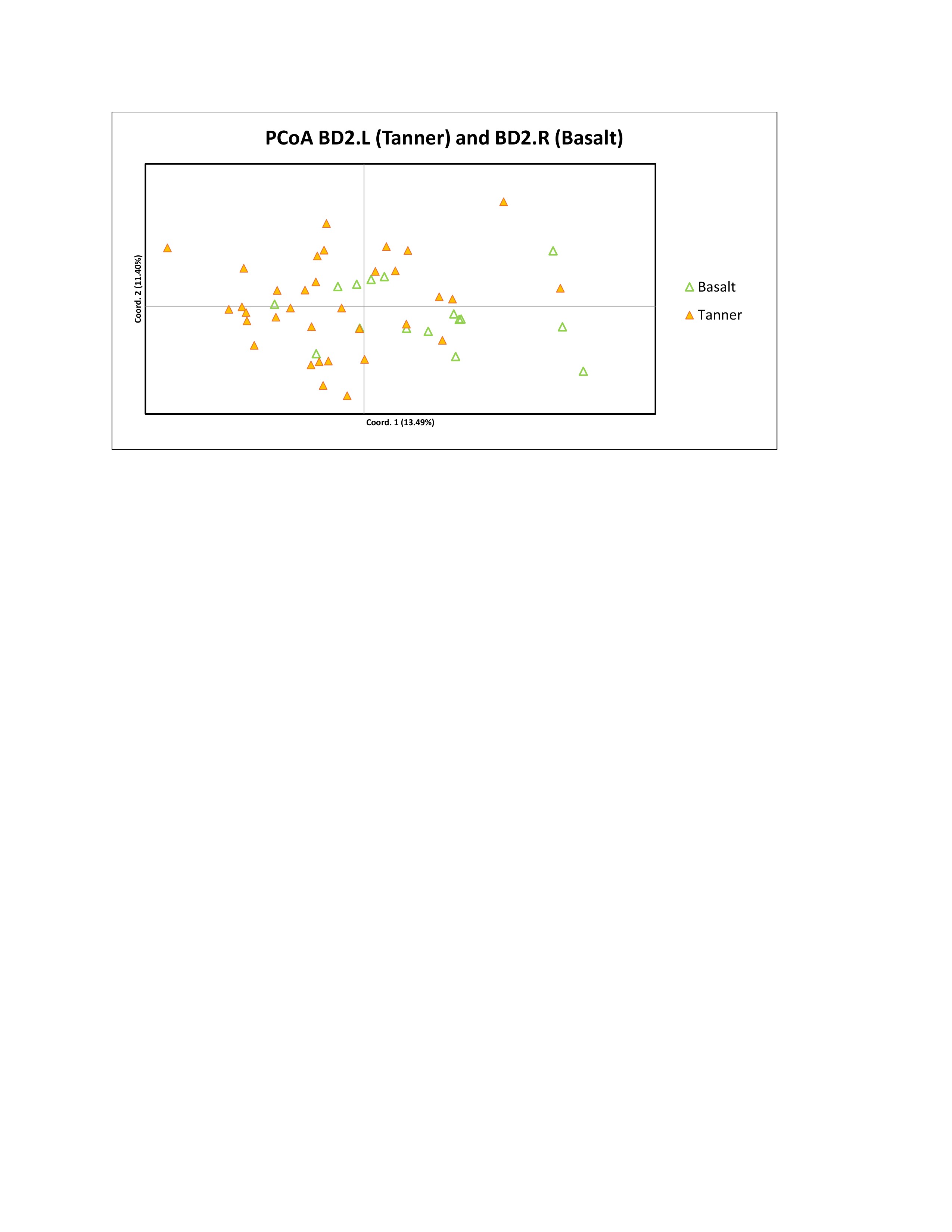

### Supplemental Figure 6B

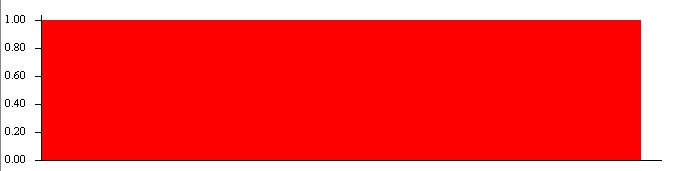
