## Supplemental Table 1 for "Is a dam-altered river in the U.S. Southwest a barrier to dispersal for populations of a common lizard, *Uta stansburiana?*"

| 8 | 241 | 6 | 64 | 58 | 16 | 34 | 20 | 49 |
|---|---|---|---|---|---|---|---|---|
| | | | BD1.R | BD1.L | BD2.R | BD2.L | AD1.R | AD1.L |
| **Individual ID** | **Pop** | **Marker1** | | **Marker2** | | **Marker3** | | **Marker4** |
| 1 | BD1.R | 230 | 230 | 190 | 192 | 316 | 324 | 171 |
| 2 | BD1.R | 230 | 230 | 192 | 196 | 324 | 324 | 171 |
| 3 | BD1.R | 230 | 230 | 192 | 192 | 324 | 326 | 171 |
| 4 | BD1.R | 230 | 230 | 192 | 192 | 326 | 326 | 171 |
| 5 | BD1.R | 230 | 230 | 190 | 192 | 326 | 330 | 171 |
| 7 | BD1.R | 230 | 230 | 190 | 190 | 326 | 326 | 171 |
| 9 | BD1.R | 230 | 230 | 190 | 192 | 324 | 326 | 171 |
| 10 | BD1.R | 230 | 230 | 190 | 192 | 324 | 324 | 171 |
| 11 | BD1.R | 230 | 230 | 190 | 192 | 324 | 326 | 171 |
| 12 | BD1.R | 230 | 230 | 188 | 192 | 326 | 326 | 171 |
| 13 | BD1.R | 230 | 230 | 190 | 192 | 316 | 324 | 171 |
| 14 | BD1.R | 230 | 230 | 190 | 196 | 326 | 326 | 171 |
| 15 | BD1.R | 230 | 230 | 190 | 192 | 326 | 326 | 171 |
| 16 | BD1.R | 230 | 230 | 190 | 192 | 316 | 326 | 171 |
| 19 | BD1.R | 230 | 230 | 190 | 192 | 326 | 326 | 171 |
| 20 | BD1.R | 230 | 230 | 190 | 190 | 326 | 328 | 171 |
| 21 | BD1.R | 230 | 230 | 192 | 192 | 316 | 326 | 171 |
| 23 | BD1.R | 230 | 230 | 190 | 196 | 324 | 330 | 171 |
| 25 | BD1.R | 230 | 230 | 188 | 192 | 344 | 344 | 163 |
| 26 | BD1.R | 230 | 230 | 190 | 192 | 316 | 326 | 171 |
| 27 | BD1.R | 230 | 230 | 190 | 192 | 324 | 326 | 171 |
| 47 | BD1.R | 230 | 230 | 190 | 196 | 324 | 330 | 171 |
| 48 | BD1.R | 230 | 230 | 188 | 196 | 324 | 326 | 171 |
| 118 | BD1.R | 230 | 230 | 190 | 192 | 326 | 328 | 171 |
| 119 | BD1.R | 230 | 230 | 190 | 190 | 324 | 326 | 171 |
| 120 | BD1.R | 230 | 230 | 190 | 190 | 326 | 328 | 171 |
| 121 | BD1.R | 230 | 230 | 190 | 192 | 326 | 326 | 171 |
| 122 | BD1.R | 230 | 230 | 196 | 196 | 320 | 324 | 163 |
| 123 | BD1.R | 230 | 230 | 192 | 192 | 326 | 326 | 171 |
| 124 | BD1.R | 230 | 230 | 190 | 190 | 324 | 326 | 171 |
| 125 | BD1.R | 230 | 230 | 190 | 192 | 316 | 330 | 171 |
| 126 | BD1.R | 228 | 230 | 192 | 192 | 326 | 330 | 171 |
| 127 | BD1.R | 230 | 230 | 188 | 192 | 316 | 326 | 171 |
| 128 | BD1.R | 230 | 230 | 188 | 190 | 326 | 330 | 171 |
| 129 | BD1.R | 230 | 230 | 190 | 192 | 316 | 326 | 171 |
| 130 | BD1.R | 230 | 230 | 192 | 196 | 326 | 326 | 171 |
| 131 | BD1.R | 228 | 230 | 188 | 190 | 324 | 324 | 171 |
| 132 | BD1.R | 230 | 230 | 192 | 192 | 324 | 326 | 171 |
| 133 | BD1.R | 230 | 230 | 188 | 188 | 320 | 326 | 171 |
| 134 | BD1.R | 230 | 230 | 190 | 192 | 324 | 326 | 171 |
| 135 | BD1.R | 230 | 230 | 192 | 192 | 324 | 326 | 171 |
| 136 | BD1.R | 228 | 230 | 192 | 196 | 316 | 326 | 171 |

| | | | | | | | | |
|---|---|---|---|---|---|---|---|---|
| 137 | BD1.R | 230 | 230 | 188 | 190 | 316 | 324 | 163 |
| 138 | BD1.R | 230 | 230 | 190 | 196 | 326 | 330 | 171 |
| 139 | BD1.R | 230 | 230 | 188 | 188 | 316 | 326 | 171 |
| 140 | BD1.R | 230 | 230 | 190 | 192 | 316 | 330 | 171 |
| 141 | BD1.R | 230 | 230 | 192 | 196 | 326 | 326 | 171 |
| 143 | BD1.R | 230 | 230 | 190 | 194 | 316 | 330 | 171 |
| 144 | BD1.R | 228 | 230 | 192 | 196 | 326 | 326 | 171 |
| 145 | BD1.R | 230 | 230 | 190 | 196 | 326 | 330 | 171 |
| 146 | BD1.R | 230 | 230 | 188 | 190 | 326 | 330 | 171 |
| 147 | BD1.R | 228 | 230 | 192 | 192 | 316 | 326 | 171 |
| 148 | BD1.R | 228 | 230 | 190 | 196 | 320 | 326 | 171 |
| 149 | BD1.R | 230 | 230 | 188 | 196 | 316 | 324 | 171 |
| 151 | BD1.R | 230 | 230 | 192 | 192 | 316 | 324 | 171 |
| 152 | BD1.R | 230 | 230 | 190 | 196 | 316 | 326 | 171 |
| 154 | BD1.R | 230 | 230 | 190 | 192 | 324 | 326 | 171 |
| 155 | BD1.R | 228 | 230 | 190 | 192 | 324 | 326 | 163 |
| 156 | BD1.R | 228 | 230 | 188 | 190 | 316 | 326 | 171 |
| 157 | BD1.R | 228 | 230 | 190 | 192 | 324 | 326 | 171 |
| 158 | BD1.R | 228 | 230 | 190 | 190 | 326 | 326 | 171 |
| 159 | BD1.R | 230 | 230 | 190 | 192 | 324 | 330 | 171 |
| 160 | BD1.R | 230 | 230 | 190 | 192 | 316 | 326 | 171 |
| 161 | BD1.R | 228 | 230 | 190 | 192 | 320 | 326 | 171 |
| 29 | BD1.L | 230 | 230 | 190 | 190 | 326 | 336 | 171 |
| 30 | BD1.L | 230 | 230 | 190 | 190 | 322 | 324 | 163 |
| 32 | BD1.L | 230 | 230 | 190 | 192 | 324 | 330 | 163 |
| 33 | BD1.L | 230 | 230 | 190 | 190 | 328 | 330 | 171 |
| 35 | BD1.L | 230 | 230 | 190 | 192 | 330 | 336 | 163 |
| 37 | BD1.L | 230 | 230 | 190 | 192 | 326 | 328 | 163 |
| 38 | BD1.L | 230 | 230 | 192 | 192 | 326 | 336 | 171 |
| 39 | BD1.L | 230 | 230 | 190 | 190 | 324 | 328 | 171 |
| 41 | BD1.L | 230 | 230 | 190 | 190 | 322 | 330 | 171 |
| 49 | BD1.L | 230 | 230 | 188 | 192 | 324 | 330 | 163 |
| 51 | BD1.L | 230 | 230 | 190 | 192 | 330 | 330 | 171 |
| 52 | BD1.L | 230 | 230 | 190 | 192 | 326 | 330 | 171 |
| 54 | BD1.L | 230 | 230 | 190 | 192 | 326 | 328 | 171 |
| 56 | BD1.L | 230 | 230 | 190 | 190 | 324 | 336 | 163 |
| 58 | BD1.L | 228 | 230 | 188 | 190 | 322 | 330 | 171 |
| 64 | BD1.L | 230 | 230 | 190 | 190 | 324 | 324 | 163 |
| 67 | BD1.L | 230 | 230 | 190 | 190 | 330 | 336 | 163 |
| 70 | BD1.L | 230 | 230 | 190 | 190 | 324 | 336 | 171 |
| 75 | BD1.L | 230 | 230 | 190 | 192 | 324 | 324 | 171 |
| 77 | BD1.L | 230 | 230 | 190 | 190 | 324 | 324 | 163 |
| 78 | BD1.L | 230 | 230 | 190 | 192 | 322 | 324 | 171 |
| 79 | BD1.L | 230 | 230 | 190 | 190 | 324 | 330 | 163 |
| 81 | BD1.L | 230 | 230 | 190 | 192 | 322 | 336 | 171 |

| | | | | | | | | |
|---|---|---|---|---|---|---|---|---|
| 82 | BD1.L | 230 | 230 | 190 | 190 | 324 | 330 | 163 |
| 84 | BD1.L | 230 | 230 | 190 | 192 | 322 | 324 | 163 |
| 85 | BD1.L | 230 | 230 | 190 | 190 | 326 | 330 | 171 |
| 86 | BD1.L | 228 | 230 | 190 | 190 | 324 | 330 | 163 |
| 87 | BD1.L | 228 | 230 | 190 | 190 | 324 | 326 | 171 |
| 88 | BD1.L | 230 | 230 | 190 | 190 | 324 | 328 | 171 |
| 89 | BD1.L | 228 | 230 | 190 | 190 | 324 | 326 | 163 |
| 90 | BD1.L | 230 | 230 | 190 | 192 | 324 | 328 | 163 |
| 91 | BD1.L | 230 | 230 | 190 | 192 | 328 | 330 | 163 |
| 92 | BD1.L | 230 | 230 | 192 | 192 | 324 | 324 | 163 |
| 93 | BD1.L | 230 | 230 | 190 | 190 | 324 | 336 | 171 |
| 94 | BD1.L | 228 | 230 | 190 | 190 | 324 | 326 | 171 |
| 95 | BD1.L | 230 | 230 | 190 | 190 | 322 | 326 | 163 |
| 96 | BD1.L | 228 | 230 | 190 | 190 | 326 | 332 | 171 |
| 97 | BD1.L | 228 | 230 | 190 | 190 | 324 | 336 | 163 |
| 98 | BD1.L | 230 | 230 | 190 | 190 | 322 | 336 | 163 |
| 99 | BD1.L | 230 | 230 | 190 | 190 | 324 | 336 | 163 |
| 100 | BD1.L | 230 | 230 | 190 | 192 | 324 | 328 | 163 |
| 101 | BD1.L | 230 | 230 | 190 | 190 | 324 | 336 | 171 |
| 102 | BD1.L | 230 | 230 | 190 | 192 | 326 | 330 | 163 |
| 103 | BD1.L | 230 | 230 | 190 | 190 | 324 | 336 | 163 |
| 104 | BD1.L | 230 | 230 | 190 | 190 | 328 | 330 | 171 |
| 105 | BD1.L | 230 | 230 | 190 | 190 | 326 | 328 | 163 |
| 106 | BD1.L | 228 | 230 | 190 | 192 | 324 | 324 | 171 |
| 107 | BD1.L | 230 | 230 | 190 | 190 | 324 | 336 | 163 |
| 108 | BD1.L | 230 | 230 | 190 | 190 | 322 | 328 | 163 |
| 109 | BD1.L | 228 | 230 | 190 | 190 | 328 | 330 | 163 |
| 110 | BD1.L | 230 | 230 | 190 | 190 | 330 | 330 | 171 |
| 111 | BD1.L | 230 | 230 | 190 | 192 | 328 | 336 | 163 |
| 112 | BD1.L | 230 | 230 | 190 | 190 | 322 | 336 | 171 |
| 113 | BD1.L | 230 | 230 | 190 | 192 | 328 | 336 | 163 |
| 114 | BD1.L | 230 | 230 | 192 | 192 | 326 | 328 | 171 |
| 115 | BD1.L | 230 | 230 | 190 | 190 | 324 | 330 | 163 |
| 116 | BD1.L | 230 | 230 | 190 | 190 | 328 | 336 | 171 |
| 117 | BD1.L | 230 | 230 | 190 | 192 | 324 | 332 | 163 |
| 196 | BD2.R | 230 | 230 | 186 | 188 | 322 | 322 | 171 |
| 197 | BD2.R | 230 | 230 | 188 | 188 | 322 | 324 | 171 |
| 198 | BD2.R | 230 | 230 | 188 | 188 | 322 | 324 | 171 |
| 199 | BD2.R | 230 | 230 | 188 | 188 | 324 | 324 | 171 |
| 200 | BD2.R | 230 | 230 | 188 | 188 | 324 | 324 | 171 |
| 201 | BD2.R | 230 | 230 | 188 | 188 | 324 | 324 | 171 |
| 202 | BD2.R | 230 | 230 | 186 | 188 | 324 | 324 | 171 |
| 203 | BD2.R | 230 | 230 | 188 | 188 | 322 | 324 | 171 |
| 204 | BD2.R | 230 | 230 | 188 | 188 | 322 | 326 | 171 |
| 205 | BD2.R | 230 | 230 | 188 | 188 | 324 | 324 | 171 |

| | | | | | | | | |
|---|---|---|---|---|---|---|---|---|
| 206 | BD2.R | 230 | 230 | 188 | 188 | 322 | 324 | 171 |
| 207 | BD2.R | 230 | 230 | 188 | 188 | 322 | 324 | 171 |
| 208 | BD2.R | 230 | 230 | 188 | 188 | 324 | 324 | 171 |
| 209 | BD2.R | 230 | 230 | 186 | 188 | 322 | 324 | 171 |
| 210 | BD2.R | 230 | 230 | 186 | 188 | 322 | 322 | 171 |
| 211 | BD2.R | 230 | 230 | 186 | 188 | 324 | 324 | 171 |
| 162 | BD2.L | 230 | 230 | 188 | 190 | 324 | 324 | 171 |
| 163 | BD2.L | 230 | 230 | 188 | 188 | 324 | 324 | 171 |
| 164 | BD2.L | 230 | 230 | 188 | 190 | 324 | 324 | 171 |
| 165 | BD2.L | 230 | 230 | 188 | 188 | 322 | 324 | 171 |
| 166 | BD2.L | 230 | 230 | 188 | 190 | 322 | 324 | 171 |
| 167 | BD2.L | 230 | 230 | 188 | 188 | 320 | 324 | 171 |
| 168 | BD2.L | 230 | 230 | 188 | 188 | 322 | 322 | 171 |
| 169 | BD2.L | 230 | 230 | 188 | 188 | 324 | 324 | 171 |
| 170 | BD2.L | 230 | 230 | 188 | 188 | 320 | 324 | 171 |
| 171 | BD2.L | 230 | 230 | 188 | 188 | 324 | 324 | 171 |
| 172 | BD2.L | 230 | 230 | 188 | 188 | 320 | 324 | 171 |
| 173 | BD2.L | 230 | 230 | 188 | 188 | 322 | 324 | 171 |
| 174 | BD2.L | 230 | 230 | 180 | 188 | 324 | 326 | 171 |
| 175 | BD2.L | 230 | 230 | 188 | 188 | 324 | 324 | 171 |
| 176 | BD2.L | 230 | 230 | 188 | 188 | 324 | 326 | 171 |
| 177 | BD2.L | 230 | 230 | 188 | 188 | 324 | 324 | 171 |
| 178 | BD2.L | 230 | 230 | 188 | 188 | 324 | 324 | 171 |
| 179 | BD2.L | 230 | 230 | 188 | 188 | 324 | 324 | 171 |
| 180 | BD2.L | 230 | 230 | 188 | 188 | 322 | 324 | 171 |
| 181 | BD2.L | 230 | 230 | 188 | 188 | 324 | 324 | 171 |
| 182 | BD2.L | 230 | 230 | 186 | 188 | 320 | 324 | 171 |
| 183 | BD2.L | 230 | 230 | 180 | 188 | 322 | 324 | 171 |
| 184 | BD2.L | 230 | 230 | 188 | 188 | 324 | 324 | 171 |
| 185 | BD2.L | 230 | 230 | 188 | 188 | 322 | 324 | 171 |
| 186 | BD2.L | 230 | 230 | 186 | 188 | 320 | 324 | 171 |
| 187 | BD2.L | 230 | 230 | 188 | 188 | 320 | 324 | 171 |
| 188 | BD2.L | 230 | 230 | 188 | 188 | 324 | 324 | 171 |
| 189 | BD2.L | 230 | 230 | 188 | 188 | 324 | 324 | 171 |
| 190 | BD2.L | 230 | 230 | 188 | 188 | 320 | 324 | 171 |
| 191 | BD2.L | 230 | 230 | 188 | 188 | 320 | 324 | 171 |
| 192 | BD2.L | 230 | 230 | 188 | 188 | 324 | 324 | 171 |
| 193 | BD2.L | 230 | 230 | 188 | 188 | 324 | 324 | 171 |
| 194 | BD2.L | 230 | 230 | 188 | 188 | 322 | 324 | 171 |
| 195 | BD2.L | 230 | 230 | 188 | 188 | 322 | 326 | 171 |
| C1 | AD1.R | 230 | 230 | 190 | 192 | 320 | 320 | 171 |
| C2 | AD1.R | 230 | 230 | 186 | 186 | 320 | 322 | 171 |
| C5 | AD1.R | 230 | 230 | 188 | 192 | 322 | 322 | 171 |
| C6 | AD1.R | 230 | 230 | 190 | 192 | 322 | 324 | 171 |
| C7 | AD1.R | 230 | 230 | 186 | 190 | 318 | 324 | 171 |

| | | | | | | | | |
|---|---|---|---|---|---|---|---|---|
| C8 | AD1.R | 230 | 230 | 192 | 192 | 320 | 324 | 171 |
| C9 | AD1.R | 230 | 230 | 186 | 192 | 322 | 322 | 171 |
| C10 | AD1.R | 230 | 230 | 186 | 188 | 320 | 324 | 171 |
| C11 | AD1.R | 230 | 244 | 188 | 192 | 324 | 324 | 171 |
| C12 | AD1.R | 230 | 230 | 186 | 188 | 324 | 324 | 171 |
| C13 | AD1.R | 230 | 230 | 190 | 192 | 320 | 324 | 171 |
| C15 | AD1.R | 230 | 230 | 188 | 192 | 322 | 324 | 171 |
| C16 | AD1.R | 230 | 230 | 188 | 192 | 324 | 324 | 171 |
| C17 | AD1.R | 230 | 230 | 190 | 192 | 322 | 328 | 171 |
| C18 | AD1.R | 230 | 230 | 188 | 190 | 324 | 324 | 171 |
| C19 | AD1.R | 230 | 230 | 186 | 188 | 320 | 322 | 171 |
| C20 | AD1.R | 230 | 230 | 186 | 186 | 322 | 324 | 171 |
| C21 | AD1.R | 230 | 230 | 186 | 188 | 322 | 324 | 171 |
| C22 | AD1.R | 230 | 230 | 190 | 192 | 320 | 322 | 171 |
| C23 | AD1.R | 230 | 230 | 186 | 192 | 322 | 324 | 171 |
| C24 | AD1.L | 230 | 244 | 186 | 186 | 320 | 324 | 171 |
| C25 | AD1.L | 230 | 230 | 186 | 186 | 316 | 322 | 171 |
| C26 | AD1.L | 230 | 244 | 186 | 190 | 322 | 324 | 171 |
| C27 | AD1.L | 230 | 244 | 192 | 192 | 320 | 322 | 171 |
| C28 | AD1.L | 230 | 230 | 186 | 192 | 322 | 322 | 171 |
| C29 | AD1.L | 230 | 244 | 186 | 188 | 318 | 322 | 171 |
| C30 | AD1.L | 230 | 230 | 186 | 186 | 322 | 324 | 171 |
| C31 | AD1.L | 230 | 230 | 186 | 186 | 322 | 324 | 171 |
| C32 | AD1.L | 230 | 230 | 186 | 186 | 322 | 324 | 171 |
| C33 | AD1.L | 230 | 230 | 186 | 186 | 322 | 324 | 171 |
| C34 | AD1.L | 230 | 230 | 186 | 186 | 322 | 324 | 171 |
| C35 | AD1.L | 230 | 244 | 186 | 192 | 322 | 322 | 171 |
| C36 | AD1.L | 230 | 230 | 186 | 186 | 324 | 324 | 171 |
| C37 | AD1.L | 230 | 230 | 186 | 190 | 324 | 324 | 171 |
| C38 | AD1.L | 230 | 244 | 186 | 186 | 322 | 322 | 171 |
| C39 | AD1.L | 230 | 244 | 186 | 186 | 322 | 330 | 171 |
| C40 | AD1.L | 230 | 230 | 186 | 186 | 322 | 324 | 171 |
| C41 | AD1.L | 230 | 230 | 186 | 186 | 324 | 326 | 171 |
| C42 | AD1.L | 230 | 230 | 186 | 186 | 322 | 322 | 171 |
| C43 | AD1.L | 230 | 230 | 186 | 186 | 322 | 322 | 171 |
| C44 | AD1.L | 230 | 230 | 186 | 186 | 324 | 326 | 171 |
| C45 | AD1.L | 230 | 230 | 186 | 186 | 316 | 322 | 171 |
| C46 | AD1.L | 230 | 230 | 186 | 186 | 322 | 324 | 171 |
| C47 | AD1.L | 230 | 230 | 186 | 190 | 322 | 326 | 171 |
| C48 | AD1.L | 230 | 230 | 186 | 186 | 322 | 324 | 171 |
| C49 | AD1.L | 230 | 230 | 186 | 190 | 322 | 326 | 171 |
| C50 | AD1.L | 230 | 244 | 186 | 186 | 320 | 322 | 171 |
| C51 | AD1.L | 230 | 230 | 186 | 186 | 322 | 324 | 171 |
| C52 | AD1.L | 230 | 230 | 186 | 186 | 322 | 322 | 171 |
| C53 | AD1.L | 230 | 244 | 186 | 186 | 322 | 322 | 171 |

| | | | | | | | | |
|---|---|---|---|---|---|---|---|---|
| C54 | AD1.L | 230 | 230 | 186 | 190 | 316 | 326 | 171 |
| C55 | AD1.L | 230 | 230 | 186 | 192 | 320 | 322 | 171 |
| C56 | AD1.L | 230 | 230 | 186 | 190 | 324 | 330 | 171 |
| C57 | AD1.L | 230 | 244 | 186 | 190 | 316 | 324 | 171 |
| C58 | AD1.L | 230 | 230 | 186 | 186 | 316 | 320 | 171 |
| C59 | AD1.L | 230 | 230 | 186 | 186 | 322 | 322 | 171 |
| C60 | AD1.L | 230 | 230 | 186 | 186 | 322 | 324 | 171 |
| C61 | AD1.L | 230 | 230 | 186 | 186 | 316 | 322 | 171 |
| C62 | AD1.L | 230 | 230 | 186 | 186 | 322 | 322 | 171 |
| C63 | AD1.L | 230 | 230 | 186 | 186 | 322 | 324 | 171 |
| C64 | AD1.L | 230 | 244 | 186 | 186 | 316 | 322 | 171 |
| C65 | AD1.L | 230 | 230 | 186 | 190 | 322 | 324 | 171 |
| C66 | AD1.L | 230 | 230 | 186 | 186 | 316 | 322 | 171 |
| C67 | AD1.L | 230 | 230 | 186 | 192 | 322 | 324 | 171 |
| C68 | AD1.L | 230 | 230 | 186 | 186 | 322 | 322 | 171 |
| C69 | AD1.L | 230 | 244 | 186 | 190 | 322 | 324 | 171 |
| C70 | AD1.L | 230 | 230 | 186 | 186 | 320 | 322 | 171 |
| C71 | AD1.L | 230 | 230 | 186 | 186 | 322 | 326 | 171 |
| C72 | AD1.L | 230 | 230 | 186 | 186 | 322 | 326 | 171 |

1 241

| | Marker6 | | Marker7 | | Marker8 | | Marker9 | |
|---|---|---|---|---|---|---|---|---|
| 171 | 206 | 208 | 235 | 239 | 225 | 231 | 135 | 135 |
| 171 | 206 | 208 | 235 | 241 | 231 | 233 | 135 | 135 |
| 171 | 208 | 212 | 235 | 239 | 219 | 229 | 135 | 135 |
| 171 | 208 | 210 | 233 | 241 | 225 | 227 | 135 | 135 |
| 171 | 208 | 210 | 239 | 241 | 219 | 227 | 135 | 135 |
| 171 | 208 | 210 | 235 | 235 | 225 | 229 | 135 | 135 |
| 171 | 210 | 210 | 237 | 241 | 223 | 233 | 135 | 135 |
| 171 | 210 | 210 | 233 | 237 | 223 | 229 | 135 | 135 |
| 171 | 210 | 214 | 235 | 235 | 225 | 229 | 135 | 135 |
| 171 | 210 | 212 | 237 | 239 | 225 | 231 | 135 | 135 |
| 171 | 206 | 210 | 235 | 235 | 223 | 227 | 135 | 135 |
| 171 | 210 | 216 | 235 | 239 | 223 | 231 | 135 | 135 |
| 171 | 208 | 210 | 235 | 235 | 223 | 229 | 135 | 135 |
| 171 | 208 | 210 | 235 | 237 | 219 | 225 | 135 | 135 |
| 171 | 208 | 212 | 241 | 247 | 229 | 231 | 135 | 135 |
| 171 | 208 | 208 | 235 | 235 | 219 | 229 | 135 | 135 |
| 171 | 206 | 210 | 237 | 239 | 223 | 225 | 135 | 135 |
| 171 | 206 | 212 | 235 | 237 | 229 | 229 | 135 | 135 |
| 171 | 206 | 206 | 235 | 235 | 223 | 229 | 135 | 135 |
| 171 | 210 | 210 | 235 | 235 | 223 | 223 | 135 | 135 |
| 171 | 206 | 206 | 237 | 237 | 223 | 223 | 135 | 135 |
| 171 | 206 | 210 | 237 | 241 | 223 | 223 | 135 | 135 |
| 171 | 210 | 210 | 235 | 235 | 229 | 229 | 135 | 135 |
| 171 | 0 | 0 | 235 | 235 | 225 | 229 | 135 | 137 |
| 171 | 208 | 210 | 235 | 235 | 223 | 231 | 135 | 135 |
| 171 | 208 | 212 | 235 | 241 | 223 | 231 | 135 | 135 |
| 171 | 208 | 210 | 235 | 237 | 225 | 233 | 135 | 135 |
| 171 | 210 | 210 | 241 | 241 | 227 | 231 | 135 | 135 |
| 171 | 208 | 210 | 235 | 239 | 229 | 229 | 135 | 135 |
| 171 | 208 | 210 | 235 | 241 | 223 | 227 | 135 | 135 |
| 171 | 208 | 210 | 241 | 241 | 229 | 229 | 135 | 135 |
| 171 | 208 | 210 | 235 | 237 | 221 | 229 | 135 | 135 |
| 171 | 210 | 210 | 235 | 235 | 229 | 229 | 135 | 135 |
| 171 | 210 | 216 | 235 | 235 | 223 | 227 | 135 | 135 |
| 171 | 208 | 210 | 233 | 235 | 219 | 227 | 135 | 135 |
| 171 | 208 | 210 | 235 | 235 | 219 | 233 | 135 | 135 |
| 171 | 210 | 210 | 235 | 239 | 217 | 231 | 135 | 135 |
| 171 | 208 | 210 | 235 | 235 | 229 | 233 | 135 | 135 |
| 171 | 0 | 0 | 235 | 235 | 229 | 229 | 135 | 135 |
| 171 | 208 | 212 | 235 | 237 | 229 | 229 | 135 | 135 |
| 171 | 206 | 212 | 235 | 245 | 229 | 231 | 135 | 135 |
| 171 | 208 | 212 | 237 | 241 | 223 | 227 | 135 | 135 |

| | | | | | | | | |
|---|---|---|---|---|---|---|---|---|
| 171 | 208 | 210 | 235 | 235 | 227 | 231 | 135 | 135 |
| 171 | 210 | 216 | 235 | 235 | 229 | 229 | 135 | 135 |
| 171 | 208 | 210 | 235 | 235 | 223 | 231 | 135 | 135 |
| 171 | 208 | 214 | 237 | 241 | 227 | 229 | 135 | 135 |
| 171 | 206 | 208 | 235 | 241 | 223 | 227 | 135 | 135 |
| 171 | 208 | 208 | 235 | 243 | 227 | 229 | 135 | 135 |
| 171 | 208 | 212 | 235 | 237 | 227 | 227 | 135 | 135 |
| 171 | 206 | 208 | 235 | 235 | 227 | 231 | 135 | 135 |
| 171 | 210 | 212 | 235 | 235 | 229 | 231 | 135 | 135 |
| 171 | 208 | 208 | 235 | 239 | 223 | 231 | 135 | 135 |
| 171 | 208 | 212 | 235 | 235 | 223 | 229 | 135 | 135 |
| 171 | 208 | 212 | 235 | 235 | 227 | 229 | 135 | 135 |
| 171 | 210 | 210 | 235 | 239 | 223 | 225 | 135 | 135 |
| 171 | 206 | 206 | 235 | 241 | 223 | 227 | 135 | 135 |
| 171 | 208 | 212 | 235 | 235 | 227 | 229 | 135 | 135 |
| 171 | 208 | 210 | 235 | 235 | 219 | 231 | 135 | 135 |
| 171 | 210 | 210 | 235 | 235 | 223 | 229 | 135 | 135 |
| 171 | 208 | 210 | 235 | 237 | 229 | 229 | 135 | 135 |
| 171 | 210 | 216 | 235 | 239 | 219 | 231 | 135 | 135 |
| 171 | 206 | 208 | 235 | 241 | 227 | 233 | 135 | 135 |
| 171 | 208 | 208 | 235 | 235 | 223 | 227 | 135 | 135 |
| 171 | 208 | 214 | 233 | 235 | 219 | 231 | 135 | 135 |
| 171 | 210 | 212 | 237 | 241 | 227 | 229 | 135 | 135 |
| 171 | 206 | 212 | 235 | 243 | 229 | 229 | 135 | 135 |
| 171 | 212 | 214 | 241 | 241 | 229 | 233 | 135 | 135 |
| 171 | 214 | 214 | 233 | 235 | 229 | 229 | 135 | 135 |
| 171 | 210 | 212 | 235 | 241 | 223 | 233 | 135 | 135 |
| 171 | 204 | 208 | 235 | 241 | 227 | 229 | 135 | 135 |
| 171 | 208 | 214 | 233 | 241 | 227 | 229 | 135 | 135 |
| 171 | 212 | 212 | 233 | 241 | 215 | 223 | 135 | 135 |
| 171 | 212 | 212 | 237 | 243 | 227 | 229 | 135 | 135 |
| 171 | 208 | 214 | 241 | 241 | 227 | 229 | 0 | 0 |
| 171 | 204 | 212 | 235 | 241 | 229 | 229 | 135 | 135 |
| 171 | 196 | 208 | 233 | 245 | 225 | 229 | 135 | 135 |
| 171 | 210 | 212 | 233 | 235 | 225 | 233 | 135 | 135 |
| 171 | 210 | 212 | 233 | 233 | 223 | 227 | 135 | 135 |
| 171 | 194 | 212 | 233 | 245 | 223 | 225 | 135 | 135 |
| 171 | 204 | 212 | 233 | 235 | 227 | 227 | 135 | 135 |
| 171 | 212 | 212 | 241 | 241 | 223 | 227 | 135 | 135 |
| 171 | 208 | 210 | 233 | 233 | 227 | 229 | 135 | 135 |
| 171 | 194 | 212 | 235 | 241 | 227 | 229 | 135 | 135 |
| 163 | 212 | 212 | 233 | 241 | 225 | 229 | 135 | 135 |
| 171 | 212 | 212 | 241 | 241 | 223 | 229 | 135 | 135 |
| 171 | 206 | 214 | 233 | 237 | 225 | 229 | 135 | 135 |
| 171 | 0 | 0 | 235 | 235 | 223 | 229 | 135 | 135 |

| | | | | | | | | |
|---|---|---|---|---|---|---|---|---|
| 171 | 212 | 214 | 233 | 233 | 229 | 229 | 135 | 135 |
| 171 | 0 | 0 | 241 | 243 | 229 | 229 | 135 | 135 |
| 171 | 212 | 212 | 235 | 241 | 229 | 229 | 135 | 135 |
| 171 | 212 | 214 | 233 | 241 | 229 | 229 | 135 | 135 |
| 171 | 208 | 214 | 235 | 241 | 215 | 229 | 135 | 135 |
| 171 | 212 | 212 | 235 | 241 | 227 | 229 | 135 | 135 |
| 171 | 212 | 212 | 243 | 245 | 227 | 229 | 135 | 135 |
| 171 | 212 | 212 | 0 | 0 | 229 | 229 | 135 | 135 |
| 171 | 206 | 212 | 235 | 241 | 229 | 229 | 135 | 135 |
| 171 | 212 | 212 | 235 | 241 | 223 | 229 | 135 | 135 |
| 171 | 194 | 204 | 235 | 235 | 229 | 229 | 135 | 135 |
| 171 | 210 | 210 | 233 | 235 | 229 | 229 | 135 | 135 |
| 171 | 210 | 212 | 235 | 243 | 223 | 229 | 135 | 135 |
| 171 | 204 | 212 | 235 | 241 | 225 | 231 | 135 | 135 |
| 171 | 210 | 210 | 233 | 235 | 229 | 229 | 135 | 135 |
| 171 | 212 | 212 | 233 | 241 | 223 | 227 | 135 | 135 |
| 171 | 212 | 212 | 235 | 243 | 227 | 229 | 135 | 135 |
| 163 | 212 | 212 | 233 | 237 | 223 | 229 | 135 | 135 |
| 171 | 194 | 214 | 241 | 245 | 223 | 223 | 135 | 135 |
| 163 | 206 | 206 | 237 | 237 | 223 | 229 | 135 | 135 |
| 163 | 206 | 212 | 237 | 241 | 229 | 229 | 135 | 135 |
| 171 | 212 | 214 | 235 | 241 | 229 | 229 | 135 | 135 |
| 171 | 212 | 212 | 237 | 241 | 227 | 229 | 135 | 135 |
| 171 | 204 | 212 | 241 | 241 | 229 | 231 | 135 | 135 |
| 171 | 208 | 214 | 241 | 241 | 227 | 229 | 135 | 135 |
| 171 | 212 | 212 | 241 | 243 | 225 | 225 | 135 | 135 |
| 171 | 208 | 210 | 233 | 243 | 223 | 229 | 135 | 135 |
| 171 | 210 | 212 | 235 | 241 | 229 | 229 | 135 | 135 |
| 171 | 206 | 214 | 235 | 241 | 227 | 231 | 135 | 137 |
| 171 | 206 | 212 | 241 | 241 | 227 | 229 | 135 | 135 |
| 171 | 206 | 214 | 235 | 241 | 227 | 231 | 135 | 135 |
| 171 | 210 | 214 | 241 | 243 | 225 | 229 | 135 | 135 |
| 171 | 212 | 212 | 233 | 243 | 227 | 227 | 135 | 135 |
| 171 | 204 | 212 | 241 | 241 | 227 | 229 | 135 | 135 |
| 163 | 206 | 212 | 241 | 243 | 227 | 229 | 135 | 135 |
| 171 | 208 | 212 | 233 | 245 | 217 | 231 | 135 | 135 |
| 171 | 204 | 214 | 233 | 243 | 231 | 237 | 135 | 135 |
| 171 | 208 | 208 | 233 | 243 | 225 | 231 | 135 | 135 |
| 171 | 206 | 206 | 233 | 243 | 229 | 231 | 135 | 135 |
| 171 | 204 | 210 | 233 | 241 | 231 | 231 | 135 | 135 |
| 171 | 206 | 208 | 243 | 245 | 211 | 231 | 135 | 135 |
| 171 | 206 | 208 | 241 | 243 | 229 | 233 | 135 | 135 |
| 179 | 208 | 210 | 233 | 243 | 227 | 237 | 135 | 135 |
| 171 | 206 | 212 | 243 | 243 | 229 | 231 | 135 | 135 |
| 171 | 204 | 204 | 233 | 243 | 225 | 229 | 135 | 135 |

| | | | | | | | | |
|---|---|---|---|---|---|---|---|---|
| 171 | 204 | 208 | 233 | 243 | 217 | 235 | 135 | 135 |
| 171 | 204 | 204 | 233 | 243 | 229 | 233 | 135 | 135 |
| 171 | 206 | 220 | 233 | 245 | 217 | 229 | 135 | 135 |
| 171 | 204 | 212 | 233 | 243 | 217 | 231 | 135 | 135 |
| 171 | 208 | 212 | 233 | 233 | 229 | 229 | 135 | 135 |
| 171 | 204 | 210 | 233 | 233 | 227 | 231 | 135 | 135 |
| 171 | 208 | 220 | 233 | 241 | 227 | 229 | 135 | 135 |
| 171 | 210 | 210 | 241 | 241 | 217 | 227 | 135 | 135 |
| 171 | 216 | 220 | 237 | 241 | 217 | 217 | 135 | 135 |
| 171 | 208 | 220 | 245 | 249 | 227 | 235 | 135 | 135 |
| 171 | 204 | 208 | 245 | 245 | 217 | 229 | 135 | 135 |
| 171 | 208 | 210 | 239 | 245 | 219 | 229 | 135 | 135 |
| 171 | 206 | 220 | 233 | 245 | 225 | 227 | 135 | 137 |
| 171 | 204 | 208 | 233 | 233 | 217 | 227 | 135 | 135 |
| 171 | 208 | 210 | 241 | 245 | 215 | 219 | 135 | 135 |
| 171 | 206 | 218 | 233 | 247 | 215 | 217 | 135 | 135 |
| 171 | 208 | 208 | 233 | 239 | 215 | 215 | 135 | 135 |
| 171 | 210 | 212 | 233 | 233 | 217 | 217 | 135 | 135 |
| 171 | 204 | 210 | 241 | 241 | 229 | 231 | 135 | 135 |
| 171 | 208 | 210 | 233 | 241 | 217 | 229 | 135 | 135 |
| 171 | 206 | 206 | 233 | 245 | 217 | 227 | 135 | 135 |
| 171 | 208 | 208 | 233 | 233 | 229 | 229 | 135 | 135 |
| 171 | 206 | 212 | 241 | 243 | 217 | 229 | 135 | 135 |
| 171 | 208 | 210 | 233 | 247 | 217 | 227 | 135 | 137 |
| 171 | 206 | 210 | 229 | 245 | 211 | 217 | 135 | 135 |
| 171 | 210 | 210 | 233 | 243 | 217 | 231 | 135 | 135 |
| 171 | 208 | 210 | 233 | 239 | 217 | 229 | 135 | 137 |
| 171 | 206 | 208 | 233 | 233 | 217 | 231 | 135 | 135 |
| 171 | 206 | 206 | 233 | 243 | 227 | 231 | 135 | 135 |
| 171 | 206 | 210 | 241 | 241 | 217 | 227 | 135 | 135 |
| 171 | 206 | 208 | 233 | 249 | 227 | 227 | 135 | 135 |
| 171 | 208 | 210 | 233 | 233 | 217 | 225 | 135 | 135 |
| 171 | 208 | 208 | 237 | 241 | 217 | 231 | 135 | 135 |
| 171 | 210 | 220 | 233 | 239 | 217 | 217 | 135 | 135 |
| 171 | 210 | 218 | 233 | 241 | 215 | 217 | 135 | 135 |
| 171 | 208 | 210 | 239 | 247 | 215 | 217 | 135 | 135 |
| 171 | 208 | 208 | 233 | 237 | 215 | 231 | 137 | 137 |
| 171 | 210 | 210 | 229 | 233 | 211 | 229 | 135 | 135 |
| 171 | 206 | 208 | 237 | 241 | 217 | 229 | 135 | 135 |
| 171 | 206 | 206 | 241 | 245 | 215 | 215 | 135 | 135 |
| 171 | 204 | 208 | 239 | 239 | 219 | 221 | 135 | 135 |
| 171 | 204 | 206 | 239 | 241 | 215 | 223 | 135 | 135 |
| 171 | 206 | 206 | 237 | 239 | 223 | 231 | 135 | 135 |
| 171 | 208 | 208 | 231 | 233 | 221 | 225 | 135 | 135 |
| 171 | 204 | 208 | 231 | 237 | 223 | 229 | 135 | 135 |

| | | | | | | | | |
|---|---|---|---|---|---|---|---|---|
| 171 | 206 | 206 | 231 | 247 | 225 | 229 | 135 | 135 |
| 171 | 206 | 208 | 237 | 239 | 225 | 231 | 135 | 135 |
| 171 | 208 | 208 | 237 | 241 | 215 | 231 | 135 | 135 |
| 171 | 206 | 208 | 237 | 241 | 229 | 231 | 135 | 135 |
| 171 | 204 | 206 | 237 | 239 | 223 | 227 | 135 | 135 |
| 171 | 204 | 206 | 237 | 239 | 223 | 229 | 135 | 135 |
| 171 | 204 | 206 | 237 | 241 | 221 | 227 | 135 | 135 |
| 171 | 206 | 208 | 237 | 241 | 221 | 221 | 135 | 135 |
| 171 | 204 | 206 | 237 | 243 | 223 | 223 | 135 | 135 |
| 171 | 204 | 208 | 239 | 241 | 221 | 231 | 135 | 135 |
| 171 | 208 | 208 | 237 | 243 | 221 | 225 | 135 | 135 |
| 171 | 204 | 208 | 237 | 239 | 221 | 225 | 135 | 135 |
| 171 | 208 | 210 | 239 | 239 | 219 | 227 | 135 | 135 |
| 171 | 206 | 206 | 239 | 243 | 223 | 231 | 135 | 135 |
| 171 | 206 | 206 | 229 | 239 | 223 | 223 | 135 | 135 |
| 171 | 202 | 208 | 239 | 239 | 213 | 217 | 135 | 137 |
| 171 | 206 | 210 | 235 | 237 | 213 | 223 | 135 | 135 |
| 171 | 206 | 210 | 237 | 237 | 217 | 217 | 135 | 135 |
| 171 | 202 | 210 | 235 | 241 | 217 | 223 | 135 | 135 |
| 171 | 204 | 210 | 235 | 241 | 217 | 217 | 135 | 135 |
| 171 | 204 | 204 | 235 | 237 | 213 | 223 | 135 | 135 |
| 171 | 210 | 214 | 237 | 241 | 217 | 223 | 135 | 135 |
| 171 | 210 | 210 | 237 | 239 | 217 | 217 | 135 | 135 |
| 171 | 210 | 212 | 239 | 239 | 227 | 227 | 135 | 135 |
| 171 | 210 | 214 | 241 | 243 | 223 | 223 | 135 | 135 |
| 171 | 206 | 208 | 233 | 239 | 217 | 225 | 135 | 135 |
| 171 | 206 | 210 | 239 | 241 | 223 | 225 | 135 | 135 |
| 171 | 204 | 210 | 243 | 243 | 223 | 223 | 135 | 135 |
| 171 | 204 | 210 | 239 | 241 | 215 | 223 | 135 | 135 |
| 171 | 206 | 214 | 235 | 239 | 217 | 223 | 135 | 135 |
| 171 | 202 | 210 | 239 | 241 | 217 | 223 | 135 | 135 |
| 171 | 204 | 204 | 239 | 239 | 217 | 223 | 135 | 135 |
| 171 | 202 | 210 | 235 | 235 | 221 | 223 | 135 | 135 |
| 171 | 208 | 210 | 239 | 241 | 213 | 225 | 135 | 135 |
| 171 | 210 | 214 | 239 | 239 | 213 | 223 | 135 | 135 |
| 171 | 202 | 208 | 235 | 239 | 215 | 217 | 135 | 135 |
| 171 | 204 | 206 | 235 | 239 | 213 | 223 | 135 | 135 |
| 171 | 210 | 210 | 235 | 241 | 217 | 225 | 135 | 135 |
| 171 | 210 | 210 | 239 | 241 | 217 | 223 | 135 | 135 |
| 171 | 202 | 208 | 239 | 241 | 223 | 223 | 135 | 135 |
| 171 | 208 | 210 | 239 | 241 | 213 | 223 | 135 | 135 |
| 171 | 210 | 214 | 239 | 239 | 213 | 235 | 135 | 135 |
| 171 | 204 | 210 | 235 | 239 | 213 | 225 | 135 | 135 |
| 171 | 210 | 210 | 239 | 239 | 219 | 225 | 135 | 135 |
| 171 | 210 | 210 | 239 | 241 | 217 | 223 | 135 | 135 |

| | | | | | | | | |
|---|---|---|---|---|---|---|---|---|
| 171 | 210 | 214 | 235 | 239 | 215 | 223 | 135 | 135 |
| 171 | 210 | 210 | 241 | 243 | 213 | 223 | 135 | 135 |
| 171 | 206 | 210 | 237 | 243 | 223 | 225 | 135 | 135 |
| 171 | 214 | 214 | 235 | 241 | 217 | 227 | 135 | 135 |
| 171 | 204 | 210 | 241 | 241 | 217 | 219 | 135 | 135 |
| 171 | 208 | 210 | 237 | 241 | 213 | 225 | 135 | 135 |
| 171 | 214 | 214 | 235 | 239 | 213 | 233 | 135 | 135 |
| 171 | 206 | 214 | 239 | 241 | 213 | 217 | 135 | 135 |
| 171 | 206 | 210 | 237 | 241 | 217 | 223 | 135 | 135 |
| 171 | 206 | 210 | 235 | 243 | 217 | 223 | 135 | 135 |
| 171 | 210 | 214 | 239 | 241 | 213 | 225 | 135 | 135 |
| 171 | 210 | 214 | 235 | 241 | 217 | 227 | 135 | 135 |
| 171 | 204 | 206 | 239 | 241 | 217 | 223 | 135 | 135 |
| 171 | 208 | 210 | 241 | 243 | 223 | 229 | 135 | 135 |
| 171 | 206 | 214 | 241 | 241 | 229 | 229 | 135 | 135 |
| 171 | 206 | 214 | 241 | 241 | 217 | 223 | 135 | 135 |
| 171 | 204 | 210 | 235 | 241 | 0 | 0 | 135 | 135 |
| 171 | 208 | 210 | 239 | 241 | 223 | 223 | 135 | 135 |
| 171 | 202 | 210 | 241 | 241 | 223 | 223 | 135 | 135 |
