## Supplemental Table 2 for "Is a dam-altered river in the U.S. Southwest a barrier to dispersal for populations of a common lizard, *Uta stansburiana?*"

**S2 Table. BAYESASS estimates of posterior mean migration rates**. Mean migration rate between pairs of populations of *Uta stansburiana* above and below the Glen Canyon Dam, AZ in 2020 and 2021, with 95% confidence intervals in parentheses. Values in bold are statistically significant.

| | Donor Population | | | | | |
|---|---|---|---|---|---|---|
| Recipient Population | BD1.R | BD1.L | BD2.L | BD2.R | AD1.R | AD1.L |
| BD1.R | 0.966 (0.938 to 0 | 0.01 (-0.008 to 0. | 0.01 (-0.005 to 0.018) | 0.01 (-0.005 to 0. | 0.01 (-0.006 to 0. | 0.01 (-0.005 to 0.016) |
| BD1.L | 0.017 (-0.012 to [illegible] | 0.96 (0.922 to 0.[illegible] | 0.01 (-0.006 to 0.02) | 0.01 (-0.006 to 0. | 0.01 (-0.006 to 0. | 0.01 (-0.005 to 0.016) |
| BD2.L | 0.009 (-0.009 to [illegible] | 0.01 (-0.009 to 0. | 0.95 (0.911 to 0.99) | 0.01 (-0.008 to 0. | 0.01 (-0.01 to 0.0 | 0.01 (-0.009 to 0.03) |
| BD2.R | 0.015 (-0.014 to [illegible] | 0.02 (-0.013 to 0. | **0.26 (0.196 to 0.316)** | 0.68 (0.651 to 0.[illegible] | 0.02 (-0.014 to 0. | 0.02 (-0.013 to 0.044) |
| AD1.R | 0.015 (-0.013 to [illegible] | 0.02 (-0.013 to 0. | 0.04 (-0.027 to 0.109) | 0.01 (-0.012 to 0. | 0.88 (0.791 to 0.[illegible] | 0.04 (-0.018 to 0.092) |
| AD1.L | 0.01 (-0.008 to 0. | 0.01 (-0.008 to 0. | 0.01 (-0.007 to 0.021) | 0.01 (-0.006 to 0. | 0.01 (-0.008 to 0. | 0.95 (0.918 to 0.989) |
