## Supplemental Table 3 for "Is a dam-altered river in the U.S. Southwest a barrier to dispersal for populations of a common lizard, *Uta stansburiana?*"

**S3 Table**. **Parentage analysis for each *Uta stansburiana* population pair.** Each parent-offspr

| 8 | | | 241 | 6 | 64 | 58 | 16 |
|---|---|---|---|---|---|---|---|
| | | | | | BD1.R | BD1.L | BD2.R |
| **Sample** | **Sex** | **Confidence** | **Site** | **10000Ms** | | **Brtt** | |
| 1 | M | | BD1.R | 230 | 230 | 190 | 192 |
| 2 | M | | BD1.R | 230 | 230 | 192 | 196 |
| 3 | M | | BD1.R | 230 | 230 | 192 | 192 |
| 4 | M | | BD1.R | 230 | 230 | 192 | 192 |
| 5 | M | | BD1.R | 230 | 230 | 190 | 192 |
| 7 | F | | BD1.R | 230 | 230 | 190 | 190 |
| 9 | F | | BD1.R | 230 | 230 | 190 | 192 |
| 10 | F | | BD1.R | 230 | 230 | 190 | 192 |
| 11 | M | | BD1.R | 230 | 230 | 190 | 192 |
| 12 | M | | BD1.R | 230 | 230 | 188 | 192 |
| 13 | M | | BD1.R | 230 | 230 | 190 | 192 |
| 14 | F | 80% | BD1.R | 230 | 230 | 190 | 196 |
| 15 | M | | BD1.R | 230 | 230 | 190 | 192 |
| 16 | F | | BD1.R | 230 | 230 | 190 | 192 |
| 19 | M | | BD1.R | 230 | 230 | 190 | 192 |
| 20 | F | | BD1.R | 230 | 230 | 190 | 190 |
| 21 | F | | BD1.R | 230 | 230 | 192 | 192 |
| 23 | M | | BD1.R | 230 | 230 | 190 | 196 |
| 25 | F | | BD1.R | 230 | 230 | 188 | 192 |
| 26 | M | | BD1.R | 230 | 230 | 190 | 192 |
| 27 | F | | BD1.R | 230 | 230 | 190 | 192 |
| 47 | F | | BD1.R | 230 | 230 | 190 | 196 |
| 48 | F | | BD1.R | 230 | 230 | 188 | 196 |
| 118 | F | | BD1.R | 230 | 230 | 190 | 192 |
| 119 | F | | BD1.R | 230 | 230 | 190 | 190 |
| 120 | F | | BD1.R | 230 | 230 | 190 | 190 |
| 121 | F | | BD1.R | 230 | 230 | 190 | 192 |
| 122 | M | | BD1.R | 230 | 230 | 196 | 196 |
| 123 | M | | BD1.R | 230 | 230 | 192 | 192 |
| 124 | M | | BD1.R | 230 | 230 | 190 | 190 |
| 125 | M | | BD1.R | 230 | 230 | 190 | 192 |
| 126 | F | | BD1.R | 228 | 230 | 192 | 192 |
| 127 | F | | BD1.R | 230 | 230 | 188 | 192 |
| 128 | M | | BD1.R | 230 | 230 | 188 | 190 |
| 129 | M | | BD1.R | 230 | 230 | 190 | 192 |
| 130 | M | | BD1.R | 230 | 230 | 192 | 196 |
| 131 | M | | BD1.R | 228 | 230 | 188 | 190 |
| 132 | F | | BD1.R | 230 | 230 | 192 | 192 |

| | | | | | | | |
|---|---|---|---|---|---|---|---|
| 133 | M | | BD1.R | 230 | 230 | 188 | 188 |
| 134 | M | | BD1.R | 230 | 230 | 190 | 192 |
| 135 | M | | BD1.R | 230 | 230 | 192 | 192 |
| 136 | F | | BD1.R | 228 | 230 | 192 | 196 |
| 137 | F | | BD1.R | 230 | 230 | 188 | 190 |
| 138 | F | | BD1.R | 230 | 230 | 190 | 196 |
| 139 | F | | BD1.R | 230 | 230 | 188 | 188 |
| 140 | F | | BD1.R | 230 | 230 | 190 | 192 |
| 141 | M | | BD1.R | 230 | 230 | 192 | 196 |
| 143 | M | | BD1.R | 230 | 230 | 190 | 194 |
| 144 | M | | BD1.R | 228 | 230 | 192 | 196 |
| 145 | M | | BD1.R | 230 | 230 | 190 | 196 |
| 146 | F | | BD1.R | 230 | 230 | 188 | 190 |
| 147 | M | | BD1.R | 228 | 230 | 192 | 192 |
| 148 | F | | BD1.R | 228 | 230 | 190 | 196 |
| 149 | F | | BD1.R | 230 | 230 | 188 | 196 |
| 151 | F | | BD1.R | 230 | 230 | 192 | 192 |
| 152 | M | | BD1.R | 230 | 230 | 190 | 196 |
| 154 | F | | BD1.R | 230 | 230 | 190 | 192 |
| 155 | M | | BD1.R | 228 | 230 | 190 | 192 |
| 156 | M | | BD1.R | 228 | 230 | 188 | 190 |
| 157 | M | | BD1.R | 228 | 230 | 190 | 192 |
| 158 | M | 80% | BD1.R | 228 | 230 | 190 | 190 |
| 159 | F | | BD1.R | 230 | 230 | 190 | 192 |
| 160 | M | | BD1.R | 230 | 230 | 190 | 192 |
| 161 | F | | BD1.R | 228 | 230 | 190 | 192 |
| 29 | M | | BD1.L | 230 | 230 | 190 | 190 |
| 30 | F | 80% | BD1.L | 230 | 230 | 190 | 190 |
| 32 | F | | BD1.L | 230 | 230 | 190 | 192 |
| 33 | F | | BD1.L | 230 | 230 | 190 | 190 |
| 35 | F | | BD1.L | 230 | 230 | 190 | 192 |
| 37 | F | | BD1.L | 230 | 230 | 190 | 192 |
| 38 | F | | BD1.L | 230 | 230 | 192 | 192 |
| 39 | F | | BD1.L | 230 | 230 | 190 | 190 |
| 41 | M | 80% | BD1.L | 230 | 230 | 190 | 190 |
| 49 | M | | BD1.L | 230 | 230 | 188 | 192 |
| 51 | F | | BD1.L | 230 | 230 | 190 | 192 |
| 52 | F | | BD1.L | 230 | 230 | 190 | 192 |
| 54 | M | | BD1.L | 230 | 230 | 190 | 192 |
| 56 | F | | BD1.L | 230 | 230 | 190 | 190 |
| 58 | M | | BD1.L | 228 | 230 | 188 | 190 |
| 64 | F | | BD1.L | 230 | 230 | 190 | 190 |
| 67 | F | | BD1.L | 230 | 230 | 190 | 190 |
| 70 | F | | BD1.L | 230 | 230 | 190 | 190 |
| 75 | F | | BD1.L | 230 | 230 | 190 | 192 |
| 77 | F | | BD1.L | 230 | 230 | 190 | 190 |

| | | | | | | | |
|---|---|---|---|---|---|---|---|
| 78 | F | | BD1.L | 230 | 230 | 190 | 192 |
| 79 | M | | BD1.L | 230 | 230 | 190 | 190 |
| 81 | M | | BD1.L | 230 | 230 | 190 | 192 |
| 82 | M | | BD1.L | 230 | 230 | 190 | 190 |
| 84 | M | | BD1.L | 230 | 230 | 190 | 192 |
| 85 | F | | BD1.L | 230 | 230 | 190 | 190 |
| 86 | M | | BD1.L | 228 | 230 | 190 | 190 |
| 87 | M | | BD1.L | 228 | 230 | 190 | 190 |
| 88 | M | | BD1.L | 230 | 230 | 190 | 190 |
| 89 | M | | BD1.L | 228 | 230 | 190 | 190 |
| 90 | M | | BD1.L | 230 | 230 | 190 | 192 |
| 91 | M | | BD1.L | 230 | 230 | 190 | 192 |
| 92 | M | | BD1.L | 230 | 230 | 192 | 192 |
| 93 | M | | BD1.L | 230 | 230 | 190 | 190 |
| 94 | F | | BD1.L | 228 | 230 | 190 | 190 |
| 95 | M | 80% | BD1.L | 230 | 230 | 190 | 190 |
| 96 | M | | BD1.L | 228 | 230 | 190 | 190 |
| 97 | F | | BD1.L | 228 | 230 | 190 | 190 |
| 98 | M | | BD1.L | 230 | 230 | 190 | 190 |
| 99 | M | | BD1.L | 230 | 230 | 190 | 190 |
| 100 | F | | BD1.L | 230 | 230 | 190 | 192 |
| 101 | F | | BD1.L | 230 | 230 | 190 | 190 |
| 102 | F | | BD1.L | 230 | 230 | 190 | 192 |
| 103 | M | | BD1.L | 230 | 230 | 190 | 190 |
| 104 | M | | BD1.L | 230 | 230 | 190 | 190 |
| 105 | M | | BD1.L | 230 | 230 | 190 | 190 |
| 106 | F | | BD1.L | 228 | 230 | 190 | 192 |
| 107 | F | | BD1.L | 230 | 230 | 190 | 190 |
| 108 | M | | BD1.L | 230 | 230 | 190 | 190 |
| 109 | M | | BD1.L | 228 | 230 | 190 | 190 |
| 110 | M | | BD1.L | 230 | 230 | 190 | 190 |
| 111 | M | | BD1.L | 230 | 230 | 190 | 192 |
| 112 | M | | BD1.L | 230 | 230 | 190 | 190 |
| 113 | M | | BD1.L | 230 | 230 | 190 | 192 |
| 114 | M | | BD1.L | 230 | 230 | 192 | 192 |
| 115 | M | | BD1.L | 230 | 230 | 190 | 190 |
| 116 | M | | BD1.L | 230 | 230 | 190 | 190 |
| 117 | F | | BD1.L | 230 | 230 | 190 | 192 |
| 196 | M | | BD2.R | 230 | 230 | 186 | 188 |
| 197 | M | | BD2.R | 230 | 230 | 188 | 188 |
| 198 | M | | BD2.R | 230 | 230 | 188 | 188 |
| 199 | F | | BD2.R | 230 | 230 | 188 | 188 |
| 200 | M | | BD2.R | 230 | 230 | 188 | 188 |
| 201 | M | | BD2.R | 230 | 230 | 188 | 188 |
| 202 | F | | BD2.R | 230 | 230 | 186 | 188 |
| 203 | F | | BD2.R | 230 | 230 | 188 | 188 |

| | | | | | | | |
|---|---|---|---|---|---|---|---|
| 204 | M | | BD2.R | 230 | 230 | 188 | 188 |
| 205 | F | 80% | BD2.R | 230 | 230 | 188 | 188 |
| 206 | M | | BD2.R | 230 | 230 | 188 | 188 |
| 207 | F | 80% | BD2.R | 230 | 230 | 188 | 188 |
| 208 | M | | BD2.R | 230 | 230 | 188 | 188 |
| 209 | M | | BD2.R | 230 | 230 | 186 | 188 |
| 210 | F | | BD2.R | 230 | 230 | 186 | 188 |
| 211 | F | | BD2.R | 230 | 230 | 186 | 188 |
| 162 | M | | BD2.L | 230 | 230 | 188 | 190 |
| 163 | M | 80% | BD2.L | 230 | 230 | 188 | 188 |
| 164 | M | | BD2.L | 230 | 230 | 188 | 190 |
| 165 | M | | BD2.L | 230 | 230 | 188 | 188 |
| 166 | M | | BD2.L | 230 | 230 | 188 | 190 |
| 167 | M | | BD2.L | 230 | 230 | 188 | 188 |
| 168 | M | | BD2.L | 230 | 230 | 188 | 188 |
| 169 | F | | BD2.L | 230 | 230 | 188 | 188 |
| 170 | F | | BD2.L | 230 | 230 | 188 | 188 |
| 171 | M | | BD2.L | 230 | 230 | 188 | 188 |
| 172 | M | 80% | BD2.L | 230 | 230 | 188 | 188 |
| 173 | M | | BD2.L | 230 | 230 | 188 | 188 |
| 174 | F | | BD2.L | 230 | 230 | 180 | 188 |
| 175 | M | | BD2.L | 230 | 230 | 188 | 188 |
| 176 | M | | BD2.L | 230 | 230 | 188 | 188 |
| 177 | M | | BD2.L | 230 | 230 | 188 | 188 |
| 178 | M | | BD2.L | 230 | 230 | 188 | 188 |
| 179 | F | | BD2.L | 230 | 230 | 188 | 188 |
| 180 | F | 95% | BD2.L | 230 | 230 | 188 | 188 |
| 181 | M | | BD2.L | 230 | 230 | 188 | 188 |
| 182 | F | | BD2.L | 230 | 230 | 186 | 188 |
| 183 | M | | BD2.L | 230 | 230 | 180 | 188 |
| 184 | M | | BD2.L | 230 | 230 | 188 | 188 |
| 185 | F | 80% | BD2.L | 230 | 230 | 188 | 188 |
| 186 | F | | BD2.L | 230 | 230 | 186 | 188 |
| 187 | M | | BD2.L | 230 | 230 | 188 | 188 |
| 188 | M | | BD2.L | 230 | 230 | 188 | 188 |
| 189 | F | | BD2.L | 230 | 230 | 188 | 188 |
| 190 | M | | BD2.L | 230 | 230 | 188 | 188 |
| 191 | M | 80% | BD2.L | 230 | 230 | 188 | 188 |
| 192 | M | | BD2.L | 230 | 230 | 188 | 188 |
| 193 | M | 95% | BD2.L | 230 | 230 | 188 | 188 |
| 194 | M | | BD2.L | 230 | 230 | 188 | 188 |
| 195 | M | | BD2.L | 230 | 230 | 188 | 188 |
| C1 | M | | AD1.R | 230 | 230 | 190 | 192 |
| C2 | F | | AD1.R | 230 | 230 | 186 | 186 |
| C5 | M | | AD1.R | 230 | 230 | 188 | 192 |
| C6 | M | | AD1.R | 230 | 230 | 190 | 192 |

| | | | | | | | |
|---|---|---|---|---|---|---|---|
| C7 | M | | AD1.R | 230 | 230 | 186 | 190 |
| C8 | M | | AD1.R | 230 | 230 | 192 | 192 |
| C9 | M | | AD1.R | 230 | 230 | 186 | 192 |
| C10 | M | | AD1.R | 230 | 230 | 186 | 188 |
| C11 | M | | AD1.R | 230 | 244 | 188 | 192 |
| C12 | M | 80% | AD1.R | 230 | 230 | 186 | 188 |
| C13 | F | | AD1.R | 230 | 230 | 190 | 192 |
| C15 | M | 80% | AD1.R | 230 | 230 | 188 | 192 |
| C16 | M | | AD1.R | 230 | 230 | 188 | 192 |
| C17 | M | 80% | AD1.R | 230 | 230 | 190 | 192 |
| C18 | M | | AD1.R | 230 | 230 | 188 | 190 |
| C19 | M | | AD1.R | 230 | 230 | 186 | 188 |
| C20 | M | | AD1.R | 230 | 230 | 186 | 186 |
| C21 | M | | AD1.R | 230 | 230 | 186 | 188 |
| C22 | F | 80% | AD1.R | 230 | 230 | 190 | 192 |
| C23 | F | | AD1.R | 230 | 230 | 186 | 192 |
| C24 | F | | AD1.L | 230 | 244 | 186 | 186 |
| C25 | M | | AD1.L | 230 | 230 | 186 | 186 |
| C26 | M | | AD1.L | 230 | 244 | 186 | 190 |
| C27 | F | | AD1.L | 230 | 244 | 192 | 192 |
| C28 | M | | AD1.L | 230 | 230 | 186 | 192 |
| C29 | F | | AD1.L | 230 | 244 | 186 | 188 |
| C30 | M | | AD1.L | 230 | 230 | 186 | 186 |
| C31 | M | | AD1.L | 230 | 230 | 186 | 186 |
| C32 | M | | AD1.L | 230 | 230 | 186 | 186 |
| C33 | F | 80% | AD1.L | 230 | 230 | 186 | 186 |
| C34 | M | | AD1.L | 230 | 230 | 186 | 186 |
| C35 | M | | AD1.L | 230 | 244 | 186 | 192 |
| C36 | F | 80% | AD1.L | 230 | 230 | 186 | 186 |
| C37 | F | | AD1.L | 230 | 230 | 186 | 190 |
| C38 | M | | AD1.L | 230 | 244 | 186 | 186 |
| C39 | M | | AD1.L | 230 | 244 | 186 | 186 |
| C40 | F | | AD1.L | 230 | 230 | 186 | 186 |
| C41 | F | | AD1.L | 230 | 230 | 186 | 186 |
| C42 | F | | AD1.L | 230 | 230 | 186 | 186 |
| C43 | F | | AD1.L | 230 | 230 | 186 | 186 |
| C44 | M | | AD1.L | 230 | 230 | 186 | 186 |
| C45 | F | | AD1.L | 230 | 230 | 186 | 186 |
| C46 | F | | AD1.L | 230 | 230 | 186 | 186 |
| C47 | M | | AD1.L | 230 | 230 | 186 | 190 |
| C48 | M | | AD1.L | 230 | 230 | 186 | 186 |
| C49 | M | | AD1.L | 230 | 230 | 186 | 190 |
| C50 | F | | AD1.L | 230 | 244 | 186 | 186 |
| C51 | M | | AD1.L | 230 | 230 | 186 | 186 |
| C52 | F | | AD1.L | 230 | 230 | 186 | 186 |
| C53 | F | | AD1.L | 230 | 244 | 186 | 186 |

| | | | | | | | |
|---|---|---|---|---|---|---|---|
| C54 | F | | AD1.L | 230 | 230 | 186 | 190 |
| C55 | M | | AD1.L | 230 | 230 | 186 | 192 |
| C56 | M | | AD1.L | 230 | 230 | 186 | 190 |
| C57 | M | 95% | AD1.L | 230 | 244 | 186 | 190 |
| C58 | M | 80% | AD1.L | 230 | 230 | 186 | 186 |
| C59 | F | | AD1.L | 230 | 230 | 186 | 186 |
| C60 | F | | AD1.L | 230 | 230 | 186 | 186 |
| C61 | M | | AD1.L | 230 | 230 | 186 | 186 |
| C62 | F | | AD1.L | 230 | 230 | 186 | 186 |
| C63 | M | | AD1.L | 230 | 230 | 186 | 186 |
| C64 | F | | AD1.L | 230 | 244 | 186 | 186 |
| C65 | F | 95% | AD1.L | 230 | 230 | 186 | 190 |
| C66 | F | 80% | AD1.L | 230 | 230 | 186 | 186 |
| C67 | M | | AD1.L | 230 | 230 | 186 | 192 |
| C68 | F | | AD1.L | 230 | 230 | 186 | 186 |
| C69 | M | | AD1.L | 230 | 244 | 186 | 190 |
| C70 | M | | AD1.L | 230 | 230 | 186 | 186 |
| C71 | F | | AD1.L | 230 | 230 | 186 | 186 |
| C72 | F | | AD1.L | 230 | 230 | 186 | 186 |

ing pair is denoted by the same color, with 80% and 95% confidence. All parent-offspring pairs were ob

| | | | | | | | | | |
|---|---|---|---|---|---|---|---|---|---|
| 34 | 20 | 49 | 2 | 100 | 141 | | | | |
| BD2.L | AD1.R | AD1.L | | | | | | | |
| **IGs** | | **MCC** | | **PKln** | | **SMcL** | | **Sphil** | |
| 316 | 324 | 171 | 171 | 206 | 208 | 235 | 239 | 225 | 231 |
| 324 | 324 | 171 | 171 | 206 | 208 | 235 | 241 | 231 | 233 |
| 324 | 326 | 171 | 171 | 208 | 212 | 235 | 239 | 219 | 229 |
| 326 | 326 | 171 | 171 | 208 | 210 | 233 | 241 | 225 | 227 |
| 326 | 330 | 171 | 171 | 208 | 210 | 239 | 241 | 219 | 227 |
| 326 | 326 | 171 | 171 | 208 | 210 | 235 | 235 | 225 | 229 |
| 324 | 326 | 171 | 171 | 210 | 210 | 237 | 241 | 223 | 233 |
| 324 | 324 | 171 | 171 | 210 | 210 | 233 | 237 | 223 | 229 |
| 324 | 326 | 171 | 171 | 210 | 214 | 235 | 235 | 225 | 229 |
| 326 | 326 | 171 | 171 | 210 | 212 | 237 | 239 | 225 | 231 |
| 316 | 324 | 171 | 171 | 206 | 210 | 235 | 235 | 223 | 227 |
| 326 | 326 | 171 | 171 | 210 | 216 | 235 | 239 | 223 | 231 |
| 326 | 326 | 171 | 171 | 208 | 210 | 235 | 235 | 223 | 229 |
| 316 | 326 | 171 | 171 | 208 | 210 | 235 | 237 | 219 | 225 |
| 326 | 326 | 171 | 171 | 208 | 212 | 241 | 247 | 229 | 231 |
| 326 | 328 | 171 | 171 | 208 | 208 | 235 | 235 | 219 | 229 |
| 316 | 326 | 171 | 171 | 206 | 210 | 237 | 239 | 223 | 225 |
| 324 | 330 | 171 | 171 | 206 | 212 | 235 | 237 | 229 | 229 |
| 344 | 344 | 163 | 171 | 206 | 206 | 235 | 235 | 223 | 229 |
| 316 | 326 | 171 | 171 | 210 | 210 | 235 | 235 | 223 | 223 |
| 324 | 326 | 171 | 171 | 206 | 206 | 237 | 237 | 223 | 223 |
| 324 | 330 | 171 | 171 | 206 | 210 | 237 | 241 | 223 | 223 |
| 324 | 326 | 171 | 171 | 210 | 210 | 235 | 235 | 229 | 229 |
| 326 | 328 | 171 | 171 | 0 | 0 | 235 | 235 | 225 | 229 |
| 324 | 326 | 171 | 171 | 208 | 210 | 235 | 235 | 223 | 231 |
| 326 | 328 | 171 | 171 | 208 | 212 | 235 | 241 | 223 | 231 |
| 326 | 326 | 171 | 171 | 208 | 210 | 235 | 237 | 225 | 233 |
| 320 | 324 | 163 | 171 | 210 | 210 | 241 | 241 | 227 | 231 |
| 326 | 326 | 171 | 171 | 208 | 210 | 235 | 239 | 229 | 229 |
| 324 | 326 | 171 | 171 | 208 | 210 | 235 | 241 | 223 | 227 |
| 316 | 330 | 171 | 171 | 208 | 210 | 241 | 241 | 229 | 229 |
| 326 | 330 | 171 | 171 | 208 | 210 | 235 | 237 | 221 | 229 |
| 316 | 326 | 171 | 171 | 210 | 210 | 235 | 235 | 229 | 229 |
| 326 | 330 | 171 | 171 | 210 | 216 | 235 | 235 | 223 | 227 |
| 316 | 326 | 171 | 171 | 208 | 210 | 233 | 235 | 219 | 227 |
| 326 | 326 | 171 | 171 | 208 | 210 | 235 | 235 | 219 | 233 |
| 324 | 324 | 171 | 171 | 210 | 210 | 235 | 239 | 217 | 231 |
| 324 | 326 | 171 | 171 | 208 | 210 | 235 | 235 | 229 | 233 |

| | | | | | | | | | |
|---|---|---|---|---|---|---|---|---|---|
| 320 | 326 | 171 | 171 | 0 | 0 | 235 | 235 | 229 | 229 |
| 324 | 326 | 171 | 171 | 208 | 212 | 235 | 237 | 229 | 229 |
| 324 | 326 | 171 | 171 | 206 | 212 | 235 | 245 | 229 | 231 |
| 316 | 326 | 171 | 171 | 208 | 212 | 237 | 241 | 223 | 227 |
| 316 | 324 | 163 | 171 | 208 | 210 | 235 | 235 | 227 | 231 |
| 326 | 330 | 171 | 171 | 210 | 216 | 235 | 235 | 229 | 229 |
| 316 | 326 | 171 | 171 | 208 | 210 | 235 | 235 | 223 | 231 |
| 316 | 330 | 171 | 171 | 208 | 214 | 237 | 241 | 227 | 229 |
| 326 | 326 | 171 | 171 | 206 | 208 | 235 | 241 | 223 | 227 |
| 316 | 330 | 171 | 171 | 208 | 208 | 235 | 243 | 227 | 229 |
| 326 | 326 | 171 | 171 | 208 | 212 | 235 | 237 | 227 | 227 |
| 326 | 330 | 171 | 171 | 206 | 208 | 235 | 235 | 227 | 231 |
| 326 | 330 | 171 | 171 | 210 | 212 | 235 | 235 | 229 | 231 |
| 316 | 326 | 171 | 171 | 208 | 208 | 235 | 239 | 223 | 231 |
| 320 | 326 | 171 | 171 | 208 | 212 | 235 | 235 | 223 | 229 |
| 316 | 324 | 171 | 171 | 208 | 212 | 235 | 235 | 227 | 229 |
| 316 | 324 | 171 | 171 | 210 | 210 | 235 | 239 | 223 | 225 |
| 316 | 326 | 171 | 171 | 206 | 206 | 235 | 241 | 223 | 227 |
| 324 | 326 | 171 | 171 | 208 | 212 | 235 | 235 | 227 | 229 |
| 324 | 326 | 163 | 171 | 208 | 210 | 235 | 235 | 219 | 231 |
| 316 | 326 | 171 | 171 | 210 | 210 | 235 | 235 | 223 | 229 |
| 324 | 326 | 171 | 171 | 208 | 210 | 235 | 237 | 229 | 229 |
| 326 | 326 | 171 | 171 | 210 | 216 | 235 | 239 | 219 | 231 |
| 324 | 330 | 171 | 171 | 206 | 208 | 235 | 241 | 227 | 233 |
| 316 | 326 | 171 | 171 | 208 | 208 | 235 | 235 | 223 | 227 |
| 320 | 326 | 171 | 171 | 208 | 214 | 233 | 235 | 219 | 231 |
| 326 | 336 | 171 | 171 | 210 | 212 | 237 | 241 | 227 | 229 |
| 322 | 324 | 163 | 171 | 206 | 212 | 235 | 243 | 229 | 229 |
| 324 | 330 | 163 | 171 | 212 | 214 | 241 | 241 | 229 | 233 |
| 328 | 330 | 171 | 171 | 214 | 214 | 233 | 235 | 229 | 229 |
| 330 | 336 | 163 | 171 | 210 | 212 | 235 | 241 | 223 | 233 |
| 326 | 328 | 163 | 171 | 204 | 208 | 235 | 241 | 227 | 229 |
| 326 | 336 | 171 | 171 | 208 | 214 | 233 | 241 | 227 | 229 |
| 324 | 328 | 171 | 171 | 212 | 212 | 233 | 241 | 215 | 223 |
| 322 | 330 | 171 | 171 | 212 | 212 | 237 | 243 | 227 | 229 |
| 324 | 330 | 163 | 171 | 208 | 214 | 241 | 241 | 227 | 229 |
| 330 | 330 | 171 | 171 | 204 | 212 | 235 | 241 | 229 | 229 |
| 326 | 330 | 171 | 171 | 196 | 208 | 233 | 245 | 225 | 229 |
| 326 | 328 | 171 | 171 | 210 | 212 | 233 | 235 | 225 | 233 |
| 324 | 336 | 163 | 171 | 210 | 212 | 233 | 233 | 223 | 227 |
| 322 | 330 | 171 | 171 | 194 | 212 | 233 | 245 | 223 | 225 |
| 324 | 324 | 163 | 171 | 204 | 212 | 233 | 235 | 227 | 227 |
| 330 | 336 | 163 | 171 | 212 | 212 | 241 | 241 | 223 | 227 |
| 324 | 336 | 171 | 171 | 208 | 210 | 233 | 233 | 227 | 229 |
| 324 | 324 | 171 | 171 | 194 | 212 | 235 | 241 | 227 | 229 |
| 324 | 324 | 163 | 163 | 212 | 212 | 233 | 241 | 225 | 229 |

| | | | | | | | | | |
|---|---|---|---|---|---|---|---|---|---|
| 322 | 324 | 171 | 171 | 212 | 212 | 241 | 241 | 223 | 229 |
| 324 | 330 | 163 | 171 | 206 | 214 | 233 | 237 | 225 | 229 |
| 322 | 336 | 171 | 171 | 0 | 0 | 235 | 235 | 223 | 229 |
| 324 | 330 | 163 | 171 | 212 | 214 | 233 | 233 | 229 | 229 |
| 322 | 324 | 163 | 171 | 0 | 0 | 241 | 243 | 229 | 229 |
| 326 | 330 | 171 | 171 | 212 | 212 | 235 | 241 | 229 | 229 |
| 324 | 330 | 163 | 171 | 212 | 214 | 233 | 241 | 229 | 229 |
| 324 | 326 | 171 | 171 | 208 | 214 | 235 | 241 | 215 | 229 |
| 324 | 328 | 171 | 171 | 212 | 212 | 235 | 241 | 227 | 229 |
| 324 | 326 | 163 | 171 | 212 | 212 | 243 | 245 | 227 | 229 |
| 324 | 328 | 163 | 171 | 212 | 212 | 0 | 0 | 229 | 229 |
| 328 | 330 | 163 | 171 | 206 | 212 | 235 | 241 | 229 | 229 |
| 324 | 324 | 163 | 171 | 212 | 212 | 235 | 241 | 223 | 229 |
| 324 | 336 | 171 | 171 | 194 | 204 | 235 | 235 | 229 | 229 |
| 324 | 326 | 171 | 171 | 210 | 210 | 233 | 235 | 229 | 229 |
| 322 | 326 | 163 | 171 | 210 | 212 | 235 | 243 | 223 | 229 |
| 326 | 332 | 171 | 171 | 204 | 212 | 235 | 241 | 225 | 231 |
| 324 | 336 | 163 | 171 | 210 | 210 | 233 | 235 | 229 | 229 |
| 322 | 336 | 163 | 171 | 212 | 212 | 233 | 241 | 223 | 227 |
| 324 | 336 | 163 | 171 | 212 | 212 | 235 | 243 | 227 | 229 |
| 324 | 328 | 163 | 163 | 212 | 212 | 233 | 237 | 223 | 229 |
| 324 | 336 | 171 | 171 | 194 | 214 | 241 | 245 | 223 | 223 |
| 326 | 330 | 163 | 163 | 206 | 206 | 237 | 237 | 223 | 229 |
| 324 | 336 | 163 | 163 | 206 | 212 | 237 | 241 | 229 | 229 |
| 328 | 330 | 171 | 171 | 212 | 214 | 235 | 241 | 229 | 229 |
| 326 | 328 | 163 | 171 | 212 | 212 | 237 | 241 | 227 | 229 |
| 324 | 324 | 171 | 171 | 204 | 212 | 241 | 241 | 229 | 231 |
| 324 | 336 | 163 | 171 | 208 | 214 | 241 | 241 | 227 | 229 |
| 322 | 328 | 163 | 171 | 212 | 212 | 241 | 243 | 225 | 225 |
| 328 | 330 | 163 | 171 | 208 | 210 | 233 | 243 | 223 | 229 |
| 330 | 330 | 171 | 171 | 210 | 212 | 235 | 241 | 229 | 229 |
| 328 | 336 | 163 | 171 | 206 | 214 | 235 | 241 | 227 | 231 |
| 322 | 336 | 171 | 171 | 206 | 212 | 241 | 241 | 227 | 229 |
| 328 | 336 | 163 | 171 | 206 | 214 | 235 | 241 | 227 | 231 |
| 326 | 328 | 171 | 171 | 210 | 214 | 241 | 243 | 225 | 229 |
| 324 | 330 | 163 | 171 | 212 | 212 | 233 | 243 | 227 | 227 |
| 328 | 336 | 171 | 171 | 204 | 212 | 241 | 241 | 227 | 229 |
| 324 | 332 | 163 | 163 | 206 | 212 | 241 | 243 | 227 | 229 |
| 322 | 322 | 171 | 171 | 208 | 212 | 233 | 245 | 217 | 231 |
| 322 | 324 | 171 | 171 | 204 | 214 | 233 | 243 | 231 | 237 |
| 322 | 324 | 171 | 171 | 208 | 208 | 233 | 243 | 225 | 231 |
| 324 | 324 | 171 | 171 | 206 | 206 | 233 | 243 | 229 | 231 |
| 324 | 324 | 171 | 171 | 204 | 210 | 233 | 241 | 231 | 231 |
| 324 | 324 | 171 | 171 | 206 | 208 | 243 | 245 | 211 | 231 |
| 324 | 324 | 171 | 171 | 206 | 208 | 241 | 243 | 229 | 233 |
| 322 | 324 | 171 | 179 | 208 | 210 | 233 | 243 | 227 | 237 |

| | | | | | | | | | |
|---|---|---|---|---|---|---|---|---|---|
| 322 | 326 | 171 | 171 | 206 | 212 | 243 | 243 | 229 | 231 |
| 324 | 324 | 171 | 171 | 204 | 204 | 233 | 243 | 225 | 229 |
| 322 | 324 | 171 | 171 | 204 | 208 | 233 | 243 | 217 | 235 |
| 322 | 324 | 171 | 171 | 204 | 204 | 233 | 243 | 229 | 233 |
| 324 | 324 | 171 | 171 | 206 | 220 | 233 | 245 | 217 | 229 |
| 322 | 324 | 171 | 171 | 204 | 212 | 233 | 243 | 217 | 231 |
| 322 | 322 | 171 | 171 | 208 | 212 | 233 | 233 | 229 | 229 |
| 324 | 324 | 171 | 171 | 204 | 210 | 233 | 233 | 227 | 231 |
| 324 | 324 | 171 | 171 | 208 | 220 | 233 | 241 | 227 | 229 |
| 324 | 324 | 171 | 171 | 210 | 210 | 241 | 241 | 217 | 227 |
| 324 | 324 | 171 | 171 | 216 | 220 | 237 | 241 | 217 | 217 |
| 322 | 324 | 171 | 171 | 208 | 220 | 245 | 249 | 227 | 235 |
| 322 | 324 | 171 | 171 | 204 | 208 | 245 | 245 | 217 | 229 |
| 320 | 324 | 171 | 171 | 208 | 210 | 239 | 245 | 219 | 229 |
| 322 | 322 | 171 | 171 | 206 | 220 | 233 | 245 | 225 | 227 |
| 324 | 324 | 171 | 171 | 204 | 208 | 233 | 233 | 217 | 227 |
| 320 | 324 | 171 | 171 | 208 | 210 | 241 | 245 | 215 | 219 |
| 324 | 324 | 171 | 171 | 206 | 218 | 233 | 247 | 215 | 217 |
| 320 | 324 | 171 | 171 | 208 | 208 | 233 | 239 | 215 | 215 |
| 322 | 324 | 171 | 171 | 210 | 212 | 233 | 233 | 217 | 217 |
| 324 | 326 | 171 | 171 | 204 | 210 | 241 | 241 | 229 | 231 |
| 324 | 324 | 171 | 171 | 208 | 210 | 233 | 241 | 217 | 229 |
| 324 | 326 | 171 | 171 | 206 | 206 | 233 | 245 | 217 | 227 |
| 324 | 324 | 171 | 171 | 208 | 208 | 233 | 233 | 229 | 229 |
| 324 | 324 | 171 | 171 | 206 | 212 | 241 | 243 | 217 | 229 |
| 324 | 324 | 171 | 171 | 208 | 210 | 233 | 247 | 217 | 227 |
| 322 | 324 | 171 | 171 | 206 | 210 | 229 | 245 | 211 | 217 |
| 324 | 324 | 171 | 171 | 210 | 210 | 233 | 243 | 217 | 231 |
| 320 | 324 | 171 | 171 | 208 | 210 | 233 | 239 | 217 | 229 |
| 322 | 324 | 171 | 171 | 206 | 208 | 233 | 233 | 217 | 231 |
| 324 | 324 | 171 | 171 | 206 | 206 | 233 | 243 | 227 | 231 |
| 322 | 324 | 171 | 171 | 206 | 210 | 241 | 241 | 217 | 227 |
| 320 | 324 | 171 | 171 | 206 | 208 | 233 | 249 | 227 | 227 |
| 320 | 324 | 171 | 171 | 208 | 210 | 233 | 233 | 217 | 225 |
| 324 | 324 | 171 | 171 | 208 | 208 | 237 | 241 | 217 | 231 |
| 324 | 324 | 171 | 171 | 210 | 220 | 233 | 239 | 217 | 217 |
| 320 | 324 | 171 | 171 | 210 | 218 | 233 | 241 | 215 | 217 |
| 320 | 324 | 171 | 171 | 208 | 210 | 239 | 247 | 215 | 217 |
| 324 | 324 | 171 | 171 | 208 | 208 | 233 | 237 | 215 | 231 |
| 324 | 324 | 171 | 171 | 210 | 210 | 229 | 233 | 211 | 229 |
| 322 | 324 | 171 | 171 | 206 | 208 | 237 | 241 | 217 | 229 |
| 322 | 326 | 171 | 171 | 206 | 206 | 241 | 245 | 215 | 215 |
| 320 | 320 | 171 | 171 | 204 | 208 | 239 | 239 | 219 | 221 |
| 320 | 322 | 171 | 171 | 204 | 206 | 239 | 241 | 215 | 223 |
| 322 | 322 | 171 | 171 | 206 | 206 | 237 | 239 | 223 | 231 |
| 322 | 324 | 171 | 171 | 208 | 208 | 231 | 233 | 221 | 225 |

| | | | | | | | | | |
|---|---|---|---|---|---|---|---|---|---|
| 318 | 324 | 171 | 171 | 204 | 208 | 231 | 237 | 223 | 229 |
| 320 | 324 | 171 | 171 | 206 | 206 | 231 | 247 | 225 | 229 |
| 322 | 322 | 171 | 171 | 206 | 208 | 237 | 239 | 225 | 231 |
| 320 | 324 | 171 | 171 | 208 | 208 | 237 | 241 | 215 | 231 |
| 324 | 324 | 171 | 171 | 206 | 208 | 237 | 241 | 229 | 231 |
| 324 | 324 | 171 | 171 | 204 | 206 | 237 | 239 | 223 | 227 |
| 320 | 324 | 171 | 171 | 204 | 206 | 237 | 239 | 223 | 229 |
| 322 | 324 | 171 | 171 | 204 | 206 | 237 | 241 | 221 | 227 |
| 324 | 324 | 171 | 171 | 206 | 208 | 237 | 241 | 221 | 221 |
| 322 | 328 | 171 | 171 | 204 | 206 | 237 | 243 | 223 | 223 |
| 324 | 324 | 171 | 171 | 204 | 208 | 239 | 241 | 221 | 231 |
| 320 | 322 | 171 | 171 | 208 | 208 | 237 | 243 | 221 | 225 |
| 322 | 324 | 171 | 171 | 204 | 208 | 237 | 239 | 221 | 225 |
| 322 | 324 | 171 | 171 | 208 | 210 | 239 | 239 | 219 | 227 |
| 320 | 322 | 171 | 171 | 206 | 206 | 239 | 243 | 223 | 231 |
| 322 | 324 | 171 | 171 | 206 | 206 | 229 | 239 | 223 | 223 |
| 320 | 324 | 171 | 171 | 202 | 208 | 239 | 239 | 213 | 217 |
| 316 | 322 | 171 | 171 | 206 | 210 | 235 | 237 | 213 | 223 |
| 322 | 324 | 171 | 171 | 206 | 210 | 237 | 237 | 217 | 217 |
| 320 | 322 | 171 | 171 | 202 | 210 | 235 | 241 | 217 | 223 |
| 322 | 322 | 171 | 171 | 204 | 210 | 235 | 241 | 217 | 217 |
| 318 | 322 | 171 | 171 | 204 | 204 | 235 | 237 | 213 | 223 |
| 322 | 324 | 171 | 171 | 210 | 214 | 237 | 241 | 217 | 223 |
| 322 | 324 | 171 | 171 | 210 | 210 | 237 | 239 | 217 | 217 |
| 322 | 324 | 171 | 171 | 210 | 212 | 239 | 239 | 227 | 227 |
| 322 | 324 | 171 | 171 | 210 | 214 | 241 | 243 | 223 | 223 |
| 322 | 324 | 171 | 171 | 206 | 208 | 233 | 239 | 217 | 225 |
| 322 | 322 | 171 | 171 | 206 | 210 | 239 | 241 | 223 | 225 |
| 324 | 324 | 171 | 171 | 204 | 210 | 243 | 243 | 223 | 223 |
| 324 | 324 | 171 | 171 | 204 | 210 | 239 | 241 | 215 | 223 |
| 322 | 322 | 171 | 171 | 206 | 214 | 235 | 239 | 217 | 223 |
| 322 | 330 | 171 | 171 | 202 | 210 | 239 | 241 | 217 | 223 |
| 322 | 324 | 171 | 171 | 204 | 204 | 239 | 239 | 217 | 223 |
| 324 | 326 | 171 | 171 | 202 | 210 | 235 | 235 | 221 | 223 |
| 322 | 322 | 171 | 171 | 208 | 210 | 239 | 241 | 213 | 225 |
| 322 | 322 | 171 | 171 | 210 | 214 | 239 | 239 | 213 | 223 |
| 324 | 326 | 171 | 171 | 202 | 208 | 235 | 239 | 215 | 217 |
| 316 | 322 | 171 | 171 | 204 | 206 | 235 | 239 | 213 | 223 |
| 322 | 324 | 171 | 171 | 210 | 210 | 235 | 241 | 217 | 225 |
| 322 | 326 | 171 | 171 | 210 | 210 | 239 | 241 | 217 | 223 |
| 322 | 324 | 171 | 171 | 202 | 208 | 239 | 241 | 223 | 223 |
| 322 | 326 | 171 | 171 | 208 | 210 | 239 | 241 | 213 | 223 |
| 320 | 322 | 171 | 171 | 210 | 214 | 239 | 239 | 213 | 235 |
| 322 | 324 | 171 | 171 | 204 | 210 | 235 | 239 | 213 | 225 |
| 322 | 322 | 171 | 171 | 210 | 210 | 239 | 239 | 219 | 225 |
| 322 | 322 | 171 | 171 | 210 | 210 | 239 | 241 | 217 | 223 |

| | | | | | | | | | |
|---|---|---|---|---|---|---|---|---|---|
| 316 | 326 | 171 | 171 | 210 | 214 | 235 | 239 | 215 | 223 |
| 320 | 322 | 171 | 171 | 210 | 210 | 241 | 243 | 213 | 223 |
| 324 | 330 | 171 | 171 | 206 | 210 | 237 | 243 | 223 | 225 |
| 316 | 324 | 171 | 171 | 214 | 214 | 235 | 241 | 217 | 227 |
| 316 | 320 | 171 | 171 | 204 | 210 | 241 | 241 | 217 | 219 |
| 322 | 322 | 171 | 171 | 208 | 210 | 237 | 241 | 213 | 225 |
| 322 | 324 | 171 | 171 | 214 | 214 | 235 | 239 | 213 | 233 |
| 316 | 322 | 171 | 171 | 206 | 214 | 239 | 241 | 213 | 217 |
| 322 | 322 | 171 | 171 | 206 | 210 | 237 | 241 | 217 | 223 |
| 322 | 324 | 171 | 171 | 206 | 210 | 235 | 243 | 217 | 223 |
| 316 | 322 | 171 | 171 | 210 | 214 | 239 | 241 | 213 | 225 |
| 322 | 324 | 171 | 171 | 210 | 214 | 235 | 241 | 217 | 227 |
| 316 | 322 | 171 | 171 | 204 | 206 | 239 | 241 | 217 | 223 |
| 322 | 324 | 171 | 171 | 208 | 210 | 241 | 243 | 223 | 229 |
| 322 | 322 | 171 | 171 | 206 | 214 | 241 | 241 | 229 | 229 |
| 322 | 324 | 171 | 171 | 206 | 214 | 241 | 241 | 217 | 223 |
| 320 | 322 | 171 | 171 | 204 | 210 | 235 | 241 | 0 | 0 |
| 322 | 326 | 171 | 171 | 208 | 210 | 239 | 241 | 223 | 223 |
| 322 | 326 | 171 | 171 | 202 | 210 | 241 | 241 | 223 | 223 |

served on the same side of the river.

**Sveg**

| | |
|---|---|
| 135 | 135 |
| 135 | 135 |
| 135 | 135 |
| 135 | 135 |
| 135 | 135 |
| 135 | 135 |
| 135 | 135 |
| 135 | 135 |
| 135 | 135 |
| 135 | 135 |
| 135 | 135 |
| 135 | 135 |
| 135 | 135 |
| 135 | 135 |
| 135 | 135 |
| 135 | 135 |
| 135 | 135 |
| 135 | 135 |
| 135 | 135 |
| 135 | 135 |
| 135 | 135 |
| 135 | 135 |
| 135 | 135 |
| 135 | 137 |
| 135 | 135 |
| 135 | 135 |
| 135 | 135 |
| 135 | 135 |
| 135 | 135 |
| 135 | 135 |
| 135 | 135 |
| 135 | 135 |
| 135 | 135 |
| 135 | 135 |
| 135 | 135 |
| 135 | 135 |
| 135 | 135 |
| 135 | 135 |

| | |
|---|---|
| 135 | 135 |
| 135 | 135 |
| 135 | 135 |
| 135 | 135 |
| 135 | 135 |
| 135 | 135 |
| 135 | 135 |
| 135 | 135 |
| 135 | 135 |
| 135 | 135 |
| 135 | 135 |
| 135 | 135 |
| 135 | 135 |
| 135 | 135 |
| 135 | 135 |
| 135 | 135 |
| 135 | 135 |
| 135 | 135 |
| 135 | 135 |
| 135 | 135 |
| 135 | 135 |
| 135 | 135 |
| 135 | 135 |
| 135 | 135 |
| 135 | 135 |
| 135 | 135 |
| 135 | 135 |
| 135 | 135 |
| 135 | 135 |
| 135 | 135 |
| 135 | 135 |
| 135 | 135 |
| 135 | 135 |
| 135 | 135 |
| 135 | 135 |
| 0 | 0 |
| 135 | 135 |
| 135 | 135 |
| 135 | 135 |
| 135 | 135 |
| 135 | 135 |
| 135 | 135 |
| 135 | 135 |
| 135 | 135 |
| 135 | 135 |
| 135 | 135 |

| | |
|---|---|
| 135 | 135 |
| 135 | 135 |
| 135 | 135 |
| 135 | 135 |
| 135 | 135 |
| 135 | 135 |
| 135 | 135 |
| 135 | 135 |
| 135 | 135 |
| 135 | 135 |
| 135 | 135 |
| 135 | 135 |
| 135 | 135 |
| 135 | 135 |
| 135 | 135 |
| 135 | 135 |
| 135 | 135 |
| 135 | 135 |
| 135 | 135 |
| 135 | 135 |
| 135 | 135 |
| 135 | 135 |
| 135 | 135 |
| 135 | 135 |
| 135 | 135 |
| 135 | 135 |
| 135 | 135 |
| 135 | 135 |
| 135 | 135 |
| 135 | 135 |
| 135 | 135 |
| 135 | 137 |
| 135 | 135 |
| 135 | 135 |
| 135 | 135 |
| 135 | 135 |
| 135 | 135 |
| 135 | 135 |
| 135 | 135 |
| 135 | 135 |
| 135 | 135 |
| 135 | 135 |
| 135 | 135 |
| 135 | 135 |
| 135 | 135 |
| 135 | 135 |

| | |
|---|---|
| 135 | 135 |
| 135 | 135 |
| 135 | 135 |
| 135 | 135 |
| 135 | 135 |
| 135 | 135 |
| 135 | 135 |
| 135 | 135 |
| 135 | 135 |
| 135 | 135 |
| 135 | 135 |
| 135 | 135 |
| 135 | 135 |
| 135 | 135 |
| 135 | 137 |
| 135 | 135 |
| 135 | 135 |
| 135 | 135 |
| 135 | 135 |
| 135 | 135 |
| 135 | 135 |
| 135 | 135 |
| 135 | 135 |
| 135 | 135 |
| 135 | 135 |
| 135 | 137 |
| 135 | 135 |
| 135 | 135 |
| 135 | 137 |
| 135 | 135 |
| 135 | 135 |
| 135 | 135 |
| 135 | 135 |
| 135 | 135 |
| 135 | 135 |
| 135 | 135 |
| 135 | 135 |
| 135 | 135 |
| 137 | 137 |
| 135 | 135 |
| 135 | 135 |
| 135 | 135 |
| 135 | 135 |
| 135 | 135 |
| 135 | 135 |
| 135 | 135 |

| | |
|---|---|
| 135 | 135 |
| 135 | 135 |
| 135 | 135 |
| 135 | 135 |
| 135 | 135 |
| 135 | 135 |
| 135 | 135 |
| 135 | 135 |
| 135 | 135 |
| 135 | 135 |
| 135 | 135 |
| 135 | 135 |
| 135 | 135 |
| 135 | 135 |
| 135 | 135 |
| 135 | 135 |
| 135 | 137 |
| 135 | 135 |
| 135 | 135 |
| 135 | 135 |
| 135 | 135 |
| 135 | 135 |
| 135 | 135 |
| 135 | 135 |
| 135 | 135 |
| 135 | 135 |
| 135 | 135 |
| 135 | 135 |
| 135 | 135 |
| 135 | 135 |
| 135 | 135 |
| 135 | 135 |
| 135 | 135 |
| 135 | 135 |
| 135 | 135 |
| 135 | 135 |
| 135 | 135 |
| 135 | 135 |
| 135 | 135 |
| 135 | 135 |
| 135 | 135 |
| 135 | 135 |
| 135 | 135 |
| 135 | 135 |
| 135 | 135 |
| 135 | 135 |

| | |
|---|---|
| 135 | 135 |
| 135 | 135 |
| 135 | 135 |
| 135 | 135 |
| 135 | 135 |
| 135 | 135 |
| 135 | 135 |
| 135 | 135 |
| 135 | 135 |
| 135 | 135 |
| 135 | 135 |
| 135 | 135 |
| 135 | 135 |
| 135 | 135 |
| 135 | 135 |
| 135 | 135 |
| 135 | 135 |
| 135 | 135 |
| 135 | 135 |
